# The genome of the plague-resistant great gerbil reveals species-specific duplication of an MHCII gene

**DOI:** 10.1101/449553

**Authors:** Pernille Nilsson, Monica H. Solbakken, Boris V. Schmid, Russell J. S. Orr, Ruichen Lv, Yujun Cui, Yajun Song, Yujiang Zhang, Nils Chr. Stenseth, Ruifu Yang, Kjetill S. Jakobsen, W. Ryan Easterday, Sissel Jentoft

## Abstract

The great gerbil (*Rhombomys opimus*) is a social rodent living in permanent, complex burrow systems distributed throughout Central Asia, where it serves as the main host of several important vector-borne infectious diseases and is defined as a key reservoir species for plague (*Yersinia pestis*). Studies from the wild have shown that the great gerbil is largely resistant to plague but the genetic basis for resistance is yet to be determined. Here, we present a highly contiguous annotated genome assembly of great gerbil, covering over 96 % of the estimated 2.47 Gb genome. Comparative genomic analyses focusing on the immune gene repertoire, reveal shared gene losses within *TLR* gene families (i.e. *TLR8, TLR10* and all members of *TLR11*-subfamily) for the Gerbillinae lineage, accompanied with signs of diversifying selection of *TLR7* and *TLR9*. Most notably, we find a great gerbil-specific duplication of the *MHCII DRB* locus. *In silico* analyses suggest that the duplicated gene provides high peptide binding affinity for *Yersiniae* epitopes. The great gerbil genome provides new insights into the genomic landscape that confers immunological resistance towards plague. The high affinity for *Yersinia* epitopes could be key in our understanding of the high resistance in great gerbils, putatively conferring a faster initiation of the adaptive immune response leading to survival of the infection. Our study demonstrates the power of studying zoonosis in natural hosts through the generation of a genome resource for further comparative and experimental work on plague survival and evolution of host-pathogen interactions.

## Main

The great gerbil (*Rhombomys opimus*) is a key plague reservoir species of Central Asia[1] whose habitat stretches from Iran to Kazakhstan to North Eastern China. This diurnal, fossorial rodent lives in arid and semi-arid deserts, and forms small family groups that reside in extensive and complex burrow systems with a large surface diameter and multiple entrances, food storage and nesting chambers [2]. Where great gerbil communities coincide with human settlements and agriculture they are often viewed as pests through the destruction of crops and as carriers of vector-borne diseases[3-5]. Great gerbil is a dominant plague host species in nearly a third of the plague reservoirs located in the vast territories of Russia, Kazakhstan and China [6].

Plague, caused by the gram-negative bacterium *Yersinia pestis*, is a common disease in wildlife rodents living in semi-arid deserts and montane steppes, as well as in tropical regions[7,8]. It is predominantly transmitted between rodents by fleas living on rodents or in rodent nests [9] and regularly spills over into human populations [10], leading to individual cases and sometimes localized plague outbreaks [11]. Historically, spillover has resulted in three major human pandemics and continues to cause annual outbreaks of human plague cases in Madagascar [12-14]. Humans have played an important role in spreading the disease globally [15]. However, they are generally dead-end hosts and the long-term persistence of plague depends on plague reservoirs, which are areas where the biotic and abiotic conditions are favoring the bacterium’s survival [5].

Most commonly plague enters the body through a subcutaneous flea-bite of an infected flea, being deposited in the dermal tissue of the skin [9,16]. Once the primary physical barriers of the mammalian immune defense have been breached, the pathogen encounters a diverse community of innate immune cells and proteins evolved to recognize and destroy invasive pathogens. Here, Toll-like receptors (TLRs) and other pattern recognition receptors (PRRs) are at the forefront and have a vital role in the recognition and initiation of immune responses. Stimulation of adaptive immunity is in turn governed by the major histocompatibility complex (MHCs). MHC class I (MHCI) and class II (MHCII) proteins present antigens to CD8+ and CD4+ T lymphocytes, respectively. In particular, the CD4+ T lymphocyte is a master activator and regulator of adaptive immune responses [17,18].

In host-pathogen interactions, both sides evolve mechanisms to overpower the other engaging in an evolutionary arms race that shapes the genetic diversity on both sides [19,20]. *Y. pestis* evoke a specialized and complex attack to evade detection and destruction by the mammalian immune system to establish infection [21]. Upon entering a mammalian host, the change in temperature to 37°C initiates a change in bacterial gene expression switching on a wealth of virulence genes whose combined action enables *Y. pestis* to evade both extracellular and intracellular immune defenses [22] at the site of infection, in the lymph node and finally in the colonized blood-rich organs [16,23-26]. The host, in addition to standard immune responses, will have to establish counter measures to overcome the *Y. pestis* strategy of suppressing and delaying the innate immune responses [27,28]. This includes recognition of pathogen, resisting the bacterial signals that induce apoptosis of antigen presenting cells (APCs) and successfully producing an inflammatory response that can overpower the infection while avoiding hyperactivation.

Like all main plague reservoir hosts great gerbils can cope remarkably well with plague infections with only a minor increase in mortality levels compared to the natural mortality (see [10,29] for details). In a laboratory setting, a very large dose of *Y. pestis* is required before a lethal dose is reached where half the injected animals die (LD50) [30]. Variation in plague resistance do exists between individual great gerbils [30] however, the genetic basis of plague resistance and the differences in survival is still unclear. The adaptive immune system requires several days to respond to an infection and *Y. pestis* progresses so rapidly that it can kill susceptible hosts within days. Consequently, the genetic background of the innate immune system could potentially play a pivotal part in plague survival and also contribute to the observed heterogeneity in plague resistance [31]. For a successful response the innate immune system would have to keep the infection in check whilst properly activating the adaptive immune system [18], which can then mount an appropriate immune response leading to a more efficient and complete clearance of the pathogen. Previous studies investigating plague resistance have indeed implicated components of both innate [32-38] and adaptive immunity [39,40]. Although, none of these studies have involved wild reservoir hosts in combination with whole-genome sequencing, an approach with increased resolution that can be used in a comparative genomic setting to investigate adaptation, evolution and disease.

The importance of studying (the genetics/genomics of) zoonosis in their natural hosts is increasingly recognized [41] and the advances in sequencing technology has made it possible and affordable to do whole-genome sequencencing of non-model species for individual and comparative analysis of hosts facing a broad range of zoonosis [41].

In this paper, we present a *de novo* whole-genome sequence assembly of the major plague host, the great gerbil. We use this new resource to investigate the genomic landscape of innate and adaptive immunity with focus on candidate genes relevant for plague resistance such as *TLRs* and MHC, through genomic comparative analyses with the closely related plague hosts Mongolian gerbil (*Meriones unguiculatus*) and sand rat (*Psammomys obesus*) and other mammals.

## Results

### Genome assembly and annotation

We sequenced the genome of a wild-caught male great gerbil, sampled from the Xinjiang Province in China, using the Illumina HiSeq 2000/2500 platform (Additional file 2: Table S1 and S2). The genome was assembled *de novo* using ALLPATHS-LG resulting in an assembly consisting of 6,390 scaffolds with an N50 of 3.6 Mb and a total size of 2.376 Gb (Table 1), thus covering 96.4 % of the estimated genome size of 2.47 Gb. Assembly assessment with CEGMA and BUSCO, which investigates the presence and completeness of conserved eukaryotic and vertebrate genes, reported 85.88 % and 87.5 % gene completeness, respectively (Table 1). We were also able to locate all 39 *HOX* genes conserved in four clusters on four separate scaffolds through gene mining (Additional file 1: Fig S1). Further genome assessment with Blobology, characterizing possible contaminations, demonstrated a low degree of contamination, reporting that more than 98.5 % of the reads/bases had top hits of Rodentia. Thus, no scaffolds were filtered from our assembly. Annotation was performed using the MAKER2 pipeline and resulted in 70 974 predicted gene models of which 22 393 protein coding genes were retained based on default filtering on Annotation Edit Distance score (AED<1).

**Table 1.** Great gerbil genome assembly statistics.

| **Assembly metrics** |  |
| --- | --- |
| Total size of scaffolds (bp) | 2 376 008 858 |
| Estimated genome size (bp) | 2 464 792 293 |
| Number of scaffolds | 6 389 |
| Scaffold N50 (bp) | 3 610 217 |
| Longest scaffold (bp) | 16 185 803 |
| Total size of contigs (bp) | 2 216 488 676 |
| Number of contigs | 106 018 |
| Contig N50 (bp) | 56 880 |
| **Assembly validation** |  |
| Complete CEGMA<sup>a</sup> genes | 85.88 % (213/248) |
| Partial CEGMA genes | 95.16 % (236/248) |
| Complete Single-copy BUSCOs<sup>b</sup> | 2 114 (69.9 %) |
| Complete duplicated BUSCOs | 21 (0.69 %) |
| Fragmented BUSCOs | 533 (17.6 %) |
| Missing BUSCOs | 377 (12.5 %) |
| Total BUSCOs searched | 3 023 |

<sup>a</sup> Based on 248 highly Conserved Eukaryotic Genes (CEGs), <sup>b</sup> Based on 3,023 vertebrate-specific
BUSCO genes
Legend: The table details scaffold and contig assembly statistics as well as results from the assembly validation on genic completeness with CEGMA and BUSCO.

### Reduced TLR repertoire in great gerbil and Gerbillinae

We characterized the entire *TLR* genetic repertoire in the great gerbil genome and found 13 *TLRs*: *TLR1-13* (Fig. 1). Of these, *TLR1-7* and *TLR9* were complete with signal peptide, ecto-domain, transmembrane domain, linker and Toll/interleukin 1 receptor (TIR) domain that phylogenetically clustered well within each respective subfamily (Table 2 and Fig.2). For the remaining five *TLRs*, we were only able to retrieve fragments of *TLR8* and *TLR10* genes and although sequences of *TLR11-13* were near full length, all three members of the *TLR11* subfamily are putative non-functional pseudogenes as they contain numerous point mutations that creates premature stop codons and frameshift-causing indels. In addition, *TLR12* contains a large deletion of 78 residues (Additional file 1: Figure S2). For *TLR8*, the recovered sequence almost exclusively covers the conserved TIR domain. Relative synteny of *TLR7* and *TLR8* on chromosome X is largely conserved in both human and published rodent genomes, as well as in the great gerbil with the fragments of *TLR8* being located upstream of the full-length sequence of *TLR7* on scaffold00186 (Additional file 1: Figure S3). The great gerbil *TLR10* fragments are located on the same scaffold as full-length *TLR1* and *TLR6* (scaffold00357), in a syntenic structure comparable to other mammals (Additional file 1: Figure S3). In addition to being far from full-length sequences, the pieces of *TLR8* and *TLR10* in the great gerbil genome have point mutations that creates premature stop codons and frameshift-causing indels (Additional file 1: Figure S2). The same TLR repertoire is seen in great gerbils’ closest relatives, Mongolian gerbil and sand rat, with near full-length sequences of *TLR12* and *TLR13* and shorter fragments of *TLR8* and *TLR10*. Interestingly, for *TLR11* only shorter fragments were located for these two species, which is in contrast to the near full-length sequence identified in great gerbil. Moreover, also in these two species premature stop codons and indel-causing frameshifts were present in both the near full-length and fragmented genes (Fig. 1 and Additional file 1: Figure S2).

**FIG. 1.**
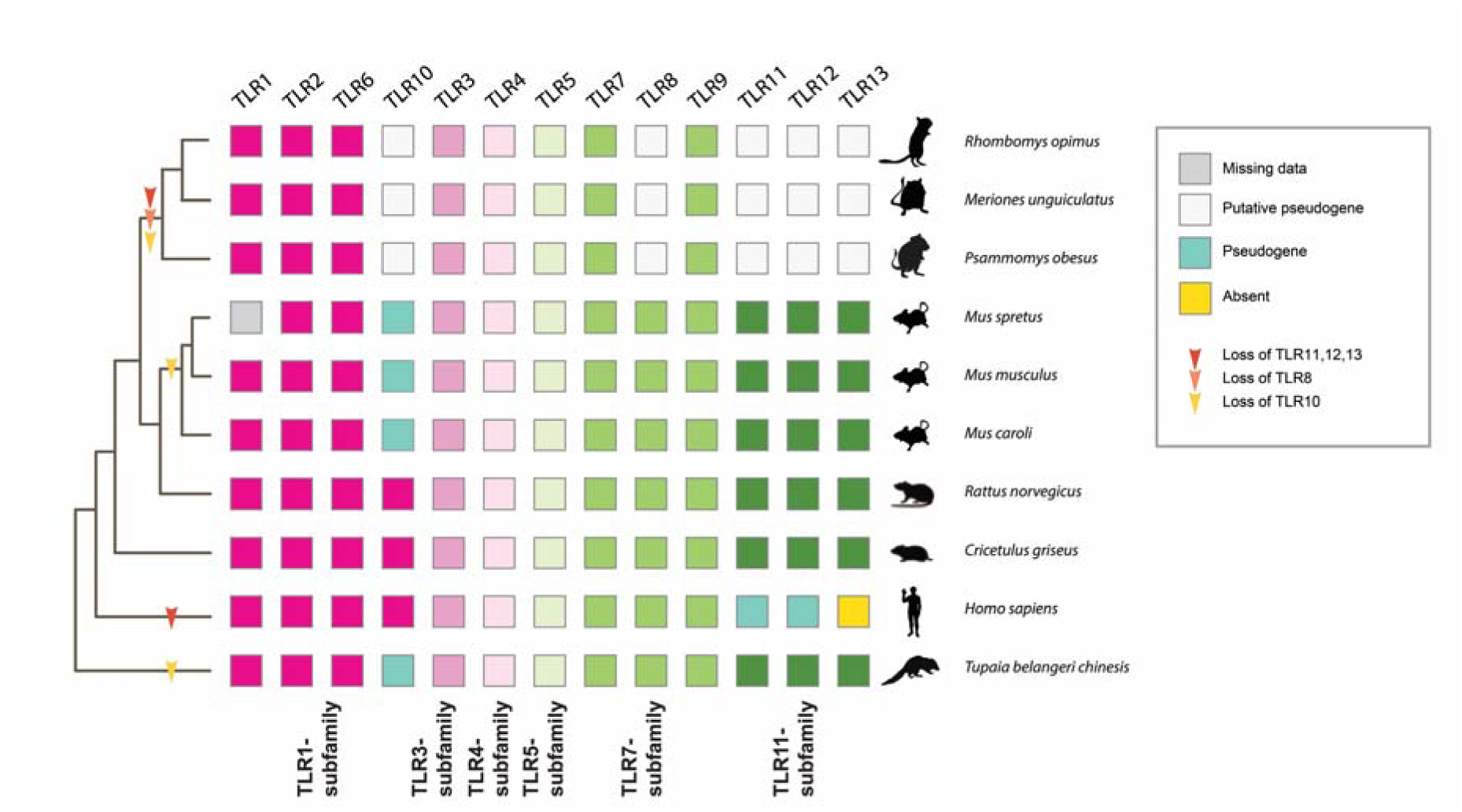
TLR repertoire in Gerbillinae compared to members of Rodentia, human and Chinese tree shrew. *TLR* repertoire of the investigated Gerbillinae, Rodentia, human and Chinese tree shrew mapped onto a composite cladogram (see Additional file 1: Figure S4). The lineage specific loss of *TLR8* and all members of the *TLR11-subfamily* in Gerbillinae and other lineage-specific *TLR* losses are marked by arrows. Depicted in boxes colored by the six major subfamilies are the individual species’ *TLR* repertoires: *TLR1-subfamily* (dark pink), *TLR3-subfamily* (pink), *TLR4-subfamily* (light pink), *TLR5-subfamily* (light green), *TLR7-subfamily* (green) and *TLR11-subfamily* (dark green). Teal colored boxes represent established pseudogenes, empty (white) boxes indicate putative pseudogenes, yellow boxes indicate complete absence of genes and grey boxes represent missing information.

**Table 2.** Overview of *TLRs* in great gerbil and their location in the genome assembly.

| Gene | Scaffold | Strand | Start | End | Size (aa) | Full-length coding sequence |
| --- | --- | --- | --- | --- | --- | --- |
| *TLR1* | scaffold00357 | + | 837 967 | 840 348 | 794 | Yes |
| *TLR2* | scaffold00513 | + | 45 595 | 47 946 | 784 | Yes |
| *TLR3* | scaffold00205 | - | 1 713 605 | 1 703 075 | 905 | Yes |
| *TLR4* | scaffold00158 | + | 3 555 106 | 3 573 424 | 838 | Yes |
| *TLR5* | scaffold00165 | - | 2 379 076 | 2 376 503 | 858 | Yes |
| *TLR6* | scaffold00357 | + | 820 802 | 823 186 | 795 | Yes |
| *TLR7* | scaffold00186 | - | 3 421 186<sup>a</sup> | 3 418 040 | 1049 | Yes<sup>b</sup> |
| *TLR8* | scaffold00186 | - |  |  | 130 | No<sup>c</sup> |
| *TLR9* | scaffold00044 | + | 7 083 426<sup>a</sup> | 7 086 521 | 1032 | Yes<sup>b</sup> |
| *TLR10* | scaffold00357 | + |  |  | 341 | No |
| *TLR11* | scaffold00001 | + |  |  | 917 | Yes<sup>d</sup> |
| *TLR12* | scaffold00071 | - | 4 008 732 | 4 006 236 | 817 | No<sup>d</sup> |
| *TLR13* | scaffold00845 | - |  |  | 899 | Yes<sup>d</sup> |

177 <sup>a</sup> Start of second codon in sequence178 <sup>b</sup> Missing start codon179 <sup>c</sup> Close to complete TIR domain plus c-terminal180 <sup>d</sup> Contains multiple point mutations and indels causing frameshifts181 Legend: The table details which scaffolds and in what orientation each *TLR* is located as well as
coordinates for the start and end of each gene (except for the pseudogenes) on the respective
scaffolds. Information on the size of the translated amino acid sequence and whether it is complete
is also shown.

**FIG. 2.**
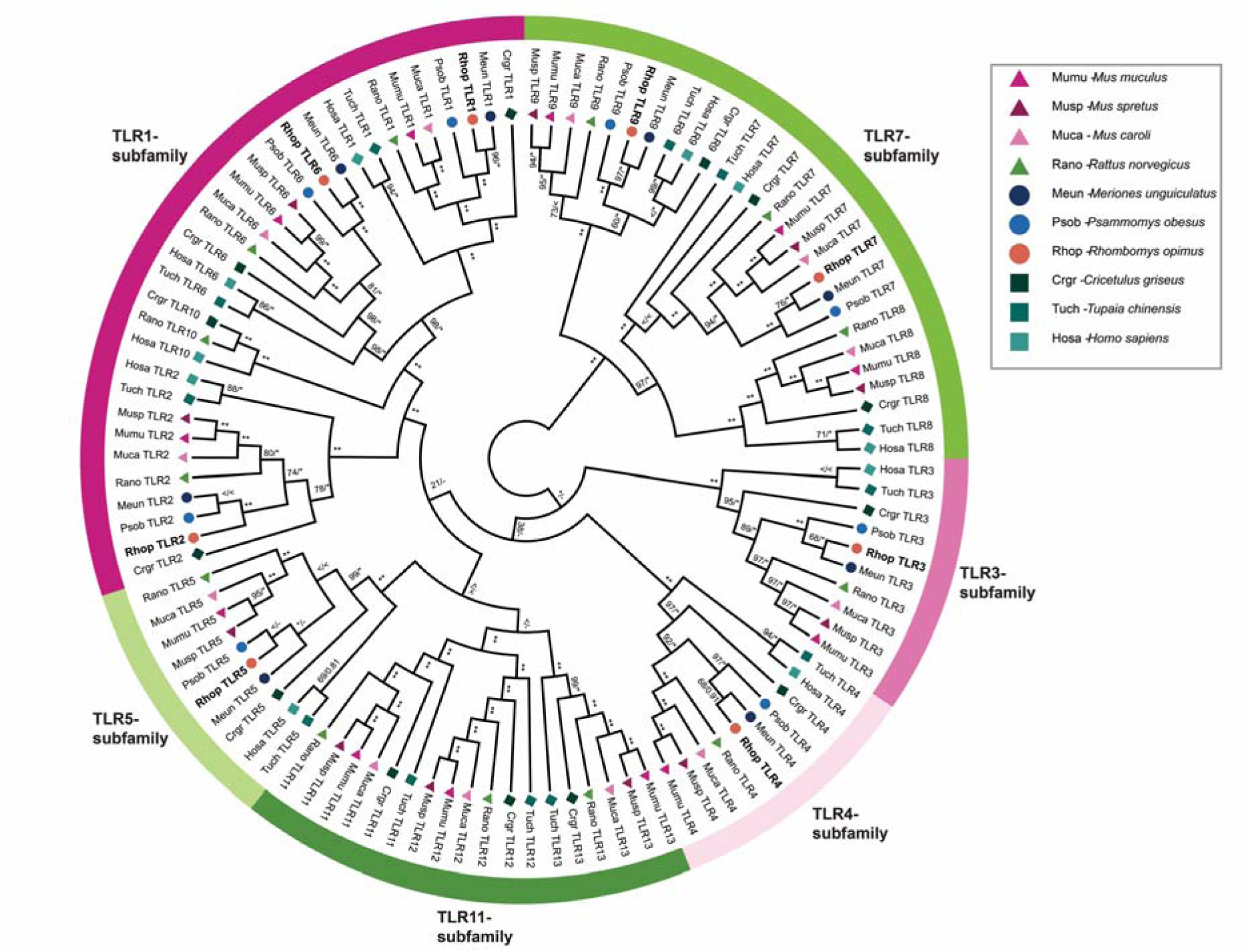
ML-phylogeny of full-length TLRs present in all investigated Gerbillinae, Rodentia, human and Chinese tree shrew. A Maximum likelihood (ML) phylogeny of nucleotide sequences all full-length *TLRs* was created using RAxML with 100x topology and 500x bootstrap replicates. A MrBayes phylogeny with 20,000,000 generations and 25 % burn-in was also created and the posterior probabilities added to the RAxML phylogeny. Great gerbil genes are marked in bold and by orange circles. The six major TLR subfamilies are marked with colored bars and corresponding names. All investigated *TLRs* including great gerbil’s, cluster well within each subfamily as well as being clearly separated into each *TLR* subfamily member.

### Diversifying selection of *TLRs*

To explore possible variations in selective pressure across the species in our analysis, we ran the adaptive branch-site random effects model (aBSREL) on all full-length *TLRs*. Evidence of episodic positive selection was demonstrated for the Gerbillinae lineage for *TLR7* and *TLR9* and for the Mongolian gerbil *TLR7* specifically (Additional file 1: Figure S5 and S6). Additionally, all full-length great gerbil *TLRs* were analyzed for sites under selection using phylogeny guided mixed effects model of evolution (MEME), from the classic datamonkey and datamonkey version 2.0 websites. Reported sites common between both analyses for all full-length *TLRs* at p-value 0.05 and their distribution among each domain of the proteins are listed in Additional file 2: Table S3. Overall, the sites under selection were almost exclusively located in the ecto-domains with a few sites located in the signal peptide (*TLR3, TLR6* and *TLR9*) and in the Linker and TIR domains (*TLR1, TLR2, TLR4* and *TLR5*). The 3D protein structure of TLR4, TLR7 and TLR9 modelled onto the human TLR5 structure further demonstrated that the sites are predominantly located in loops interspersed between the leucine-rich repeats (Additional file 1: Figures S7-9).

Scrutiny of the TLR4 amino acid sequence alignment revealed drastic differences in the properties of the residues at two positions reported to be important for maintaining signaling of hypoacetylated lipopolysaccharide (LPS). In rat (*Rattus norvegicus*) and all mouse species used in this study, the residues at position 367 and 434 are basic and positively charged while for the remaining species in the alignment including all Gerbillinae, the residues are acidic and negatively charged.

### Characterization of the great gerbil class I MHC region

The overall synteny of the MHCI region is well conserved in great gerbil, displaying the same translocation of some *MHCI* genes upstream of the MHCII region as demonstrated in mouse and rat i.e. with a distinct separation of the MHCI region into two clusters (Fig. 3). Some of the great gerbil copies were not included in the phylogeny due to missing data, which hindered their annotation. Additionally, the annotation was obstructed either by the copies being located on scaffolds not containing framework genes or due to variation in the micro-synteny of those particular loci of *MHCIa* and *MHCIb* between mouse, rat and great gerbil (Fig. 3). From the synteny it appears that *MHCI* genes are missing in the region between framework genes *Trim39* and *Trim26* and possibly between *Bat1* and *Pou5ƒ1* in the great gerbil. For full gene names for these and other framework genes mentioned below, see Additional file 2: Table S4.

**FIG. 3.**
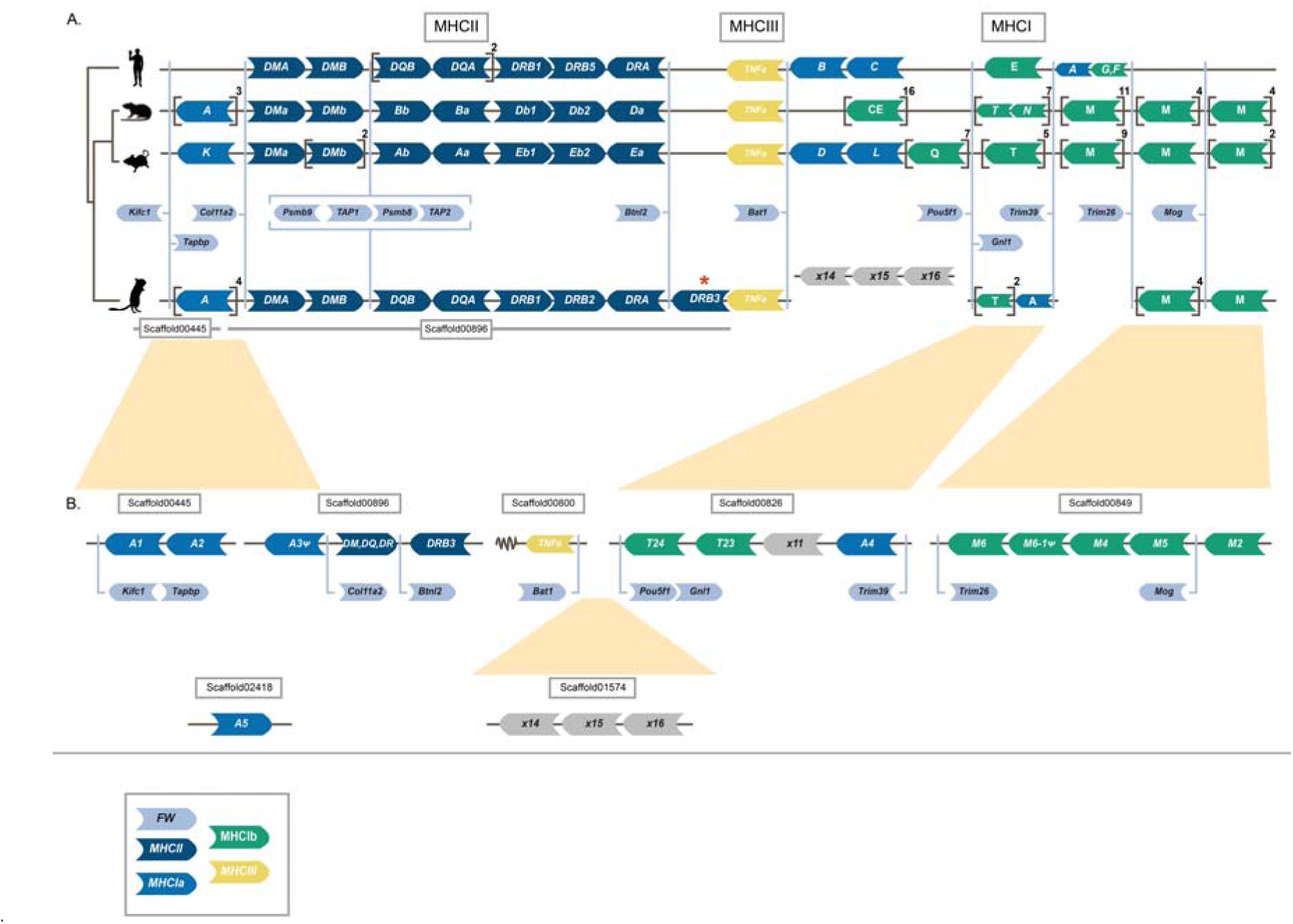
Synteny of genes in the Major histocompatibility (MHC) region of human, rat, mouse and gerbil. Genomic synteny of genes in the MHC regions of human, mouse, rat and great gerbil mapped onto a cladogram. Genes are represented by arrow-shaped boxes indicating the genomic orientation. The boxes are colored by class region and for class I by classical (Ia) or non-classical (Ib) subdivision: F ra mework (FW) genes (light blue), *MHCII* (dark blue), *MHCIa* (blue), *MHCIII* (yellow) and *MHCIb* (green). Squ are brackets indicate multiple gene copies not displayed for practical and visualization purposes, but copy number is indicated outside in superscript. Due to limitations in space and to emphasize the conserved synteny of FW genes across lineages, the genes are placed in between the general syntenies and their respective locations are indicated by light blue lines. The light blue brackets surrounding the *Psmb* and *TAP* genes indicates their constitutive organization. Putative pseudogenes are denoted with Ψ. For visualization purposes, genes of the *DP* (termed H in rat) and *DO* (termed *O* in mouse) loci are excluded. The location of all great gerbil *MHCII* genes including *Rhop-DP* and *Rhop-DO* can be found in Table 3. (A) Synteny of all MHC regions detailing *MCHI* and *II*. Panel (B) further details the genomic locations of great gerbil *MHCI* genes as indicated by the presence of FW genes located on the scaffolds and inferred from synteny comparisons with human, rat and mouse regions and phylogenetic analysis (see Fig. 4).

We were able to identify six scaffolds containing *MHCI* genes (Fig. 3 and Additional file 2: Table S5. Four of the scaffolds contained framework genes that enabled us to orient them. In total, we located 16 *MHCI* copies, of which we were able to obtain all three α domains for 10 of the copies. Three copies contain 2 out of 3 domains while for the last three copies we were only able to locate the α3 domain. In one instance, the missing α domain was due to an assembly gap. Reciprocal BLAST confirmed hits as *MHCI* genes. Due to high similarity between different *MHCI* lineages annotation of identified sequences was done through phylogenetic analyses and synteny. Our phylogeny reveals both inter-and intraspecific clustering of the great gerbil *MHCI* genes with other rodent genes with decent statistical support (i.e. bootstrap and/or posterior probabilities) of the internal branches (Fig. 4). Five great gerbil *MHCI* genes (RhopA1-5) cluster together in a main monophyletic clade while the remaining copies cluster with mouse and rat *MHCIb* genes. Two of the copies (Rhop-A3Ψ and Rhop-M6Ψ) appear to be pseudogenes as indicated by the presence of point mutations and frameshift-causing indels. Additionally, our phylogeny displays a monophyletic clustering of human *MHCI* genes (Fig. 4). The clade containing five of the great gerbil *MHCI* genes (Rhop-A1-5) possibly include a combination of both classical (*MHCIa*) and non-classical (*MHCIb*) genes as is the case for mouse and rat, where certain *MHCIb* genes cluster closely with *MHCIa* genes (Fig. 3 and Fig. 4). Also, due to the high degree of sequence similarity of rodent MHCI genes the phylogenetic relationship between clades containing non-classical M and T *MHCI* genes could not be resolved by sufficient statistical support.

**FIG. 4.**
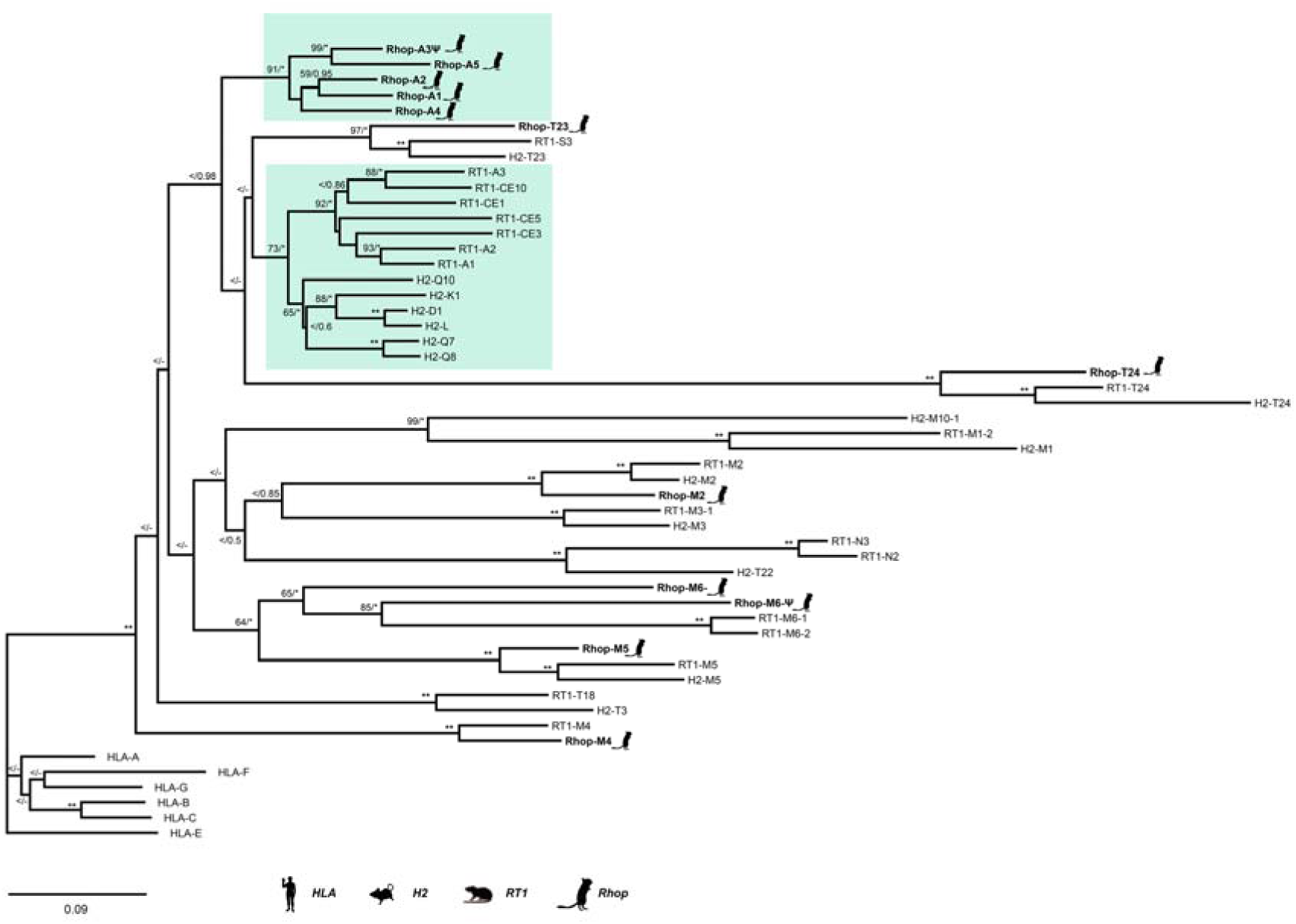
A ML-phylogeny made of nucleotide sequences of the three alpha domains from MHCI. A Maximum likelihood phylogeny of nucleotide sequences containing the three α domains of *MHCI* was created using RAxML with 100x topology and 500x bootstrap replicates. A MrBayes phylogeny with 20,000,000 generations and 25 % burn-in was also created and the posterior probabilities added to the RAxML phylogeny. BS/PP; “*” = BS 100 or PP > 0.96; “**” = BS of 100 and PP>0.97; “<” = support values below 50/0.8 and “-“ = node not present in Bayesian analysis. Twelve of the 16 great gerbil sequences were used in the analysis and are marked with a gerbil silhouette and in bold lettering. The remaining four *MHCI* sequences were excluded from the phylogenetic analyses due to missing data exceeding the set threshold of 50 %. The clusters containing *MHCIa* (classical *MHCI*) and the closest related *MHCIb* genes are marked by teal boxes. Putative pseudogenes are denoted with Ψ.

The overall synteny of the MHCI and II regions are very well conserved in great gerbil displaying the same translocation of *MHCI* genes upstream of *MHCII* as seen in mouse and rat and resulting in the separation of the *MHCI* region into two. Most notably, for *MHCII* there is a duplication of a *β* gene of the *DR* locus in great gerbil (highlighted by a red asterix) whose orientation has changed and is located downstream of the FW gene *Btnl2* that normally represents the end of the MHCII region.

### Characterization of the great gerbil class II MHC region

A single scaffold (scaffold00896) of 471 076 bp was identified to contain all genes of the MHCII region, flanked by the reference framework genes *Col11a2* and *Btnl2*. We were able to obtain orthologues of α and β genes of the classical MHCII molecules DP, DQ and DR as well as for the ‘non-classical’ DM and DO molecules (Table 3). The antigen-processing genes for the class I presentation pathway, *Psmb9*, *TAP1*, *Psmb8* and *TAP2* also maps to scaffold00896 (Fig. 3). Synteny of the MHCII region was largely conserved in great gerbil when compared to mouse, rat and human regions except for a single duplicated copy of *Rhop-DRB* (*Rhop-DRB3*) that was located distal to the *Btnl2* framework gene representing the border between class II and III of the MHC region (Fig. 3). The duplicated copy of the *Rhop-DRB* gene has an antisense orientation in contrast to the other copies of the *Rhop-DRB* genes in great gerbil. In rodents, the DR locus contains a duplication of the β gene and the two copies are termed *β*1 and *β*2, with the *β*2 gene being less polymorphic than the highly polymorphic *β*1 gene. The relative orientation of the β and α genes of the DR locus is conserved in most eutherian mammals studied to date with the genes facing each other, as is the case for Rhop-DRB1, Rhop-DRB2 and Rhop-DRA (Fig. 3). Sequence alignment and a maximum likelihood (ML) phylogeny establishes Rhop-DRB3 to be a duplication of Rhop-DRB1 (Fig. 5). Rhop-DRB1 and Rhop-DRB3 are separated by around 80 kb containing *Rhop-DRB2, Rhop-DRA* and five assembly gaps (Table 3).

**FIG. 5.**
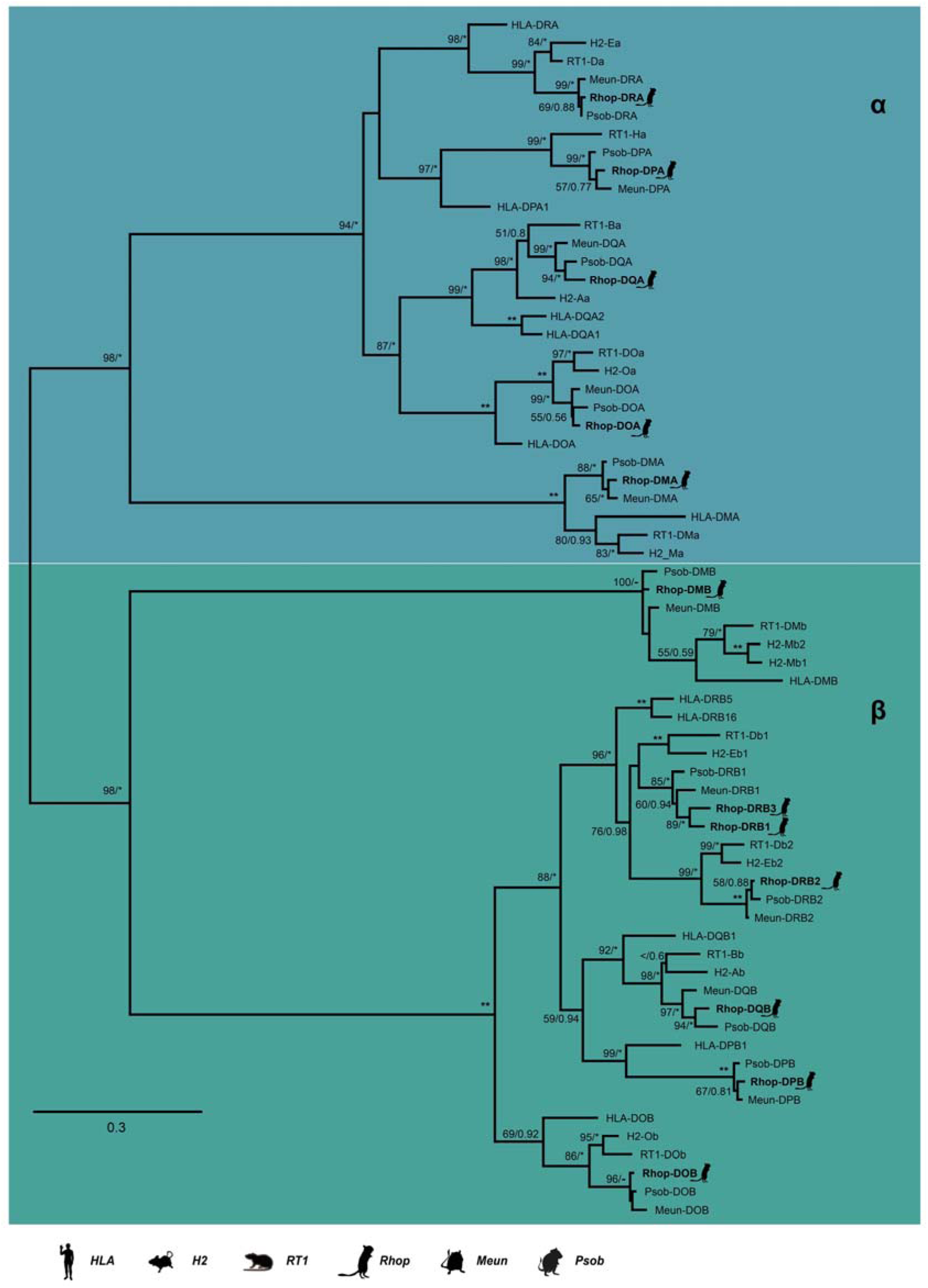
A ML-phylogeny made of nucleotide sequences of domains only from MHCII α and β genes. A Maximum likelihood phylogeny of nucleotide sequences containing the α and β domains of *MHCII α* and *β* genes was created using RAxML with 100x topology and 500x bootstrap replicates. A MrBayes phylogeny with 20,000,000 generations and 25 % burn-in was also created and the posterior probabilities added to the RAxML phylogeny. BS/PP; “*” = BS 100 or PP > 0.96; “**” = BS of 100 and PP>0.97; “<” = support values below 50/0.8 and “-” = node not present in Bayesian analysis. Great gerbil genes are indicated with bold lettering and by silhouettes. The 12 great gerbil *MHCII* genes located in the genome assembly cluster accordingly with the orthologues of human, mouse, rat, sand rat and Mongolian gerbil. The *Rhop-DRB* duplication (*Rhop-DRB3*) cluster closely with the *Rhop-DRB1* and other *DRB1* orthologs with good support. The nomenclature of *MHCII* genes in Gerbillinae are in concordance with the recommendations of the MHC Nomenclature report [43].

**Table 3.** Overview of great gerbil *MHCII* genes and their location in the genome assembly.

| Gene | Scaffold | Strand | Start | End | Size (aa) | Full-length coding sequence |
| --- | --- | --- | --- | --- | --- | --- |
| *Rhop-DPB* | scaffold00896 | - | 116 069 | 105 406 | 264 | Yes |
| *Rhop-DPA* | scaffold00896 | + | 117 514 | 120 092 | 252 | Yes |
| *Rhop-DOA* | scaffold00896 | + | 125 319 | 127 406 | 241 | Yes<sup>a</sup> |
| *Rhop-DMA* | scaffold00896 | + | 166 162 | 168 926 | 265 | Yes<sup>b</sup> |
| *Rhop-DMb* | scaffold00896 | + | 175 763 | 182 090 | 257 | Yes |
| *Rhop-DOB* | scaffold00896 | + | 239 915 | 245 006 | 172 | No<sup>c</sup> |
| *Rhop-DQB* | scaffold00896 | + | 271 007 | 278 482 | 231 | No<sup>d</sup> |
| *Rhop-DQA* | scaffold00896 | - | 294 074 | 290 301 | 255 | Yes |
| *Rhop-DRB1* | scaffold00896 | + | 310 644 | 319 368 | 265 | Yes |
| *Rhop-DRB2* | scaffold00896 | + | 326 384 | 347 931 | 272 | Yes<sup>e</sup> |
| *Rhop-DRA* | scaffold00896 | - | 355 351 | 351 425 | 254 | Yes |
| *Rhop-DRB3* | scaffold00896 | - | 403 768 | 395 068 | 265 | Yes |

<sup>a</sup> Missing final residue and stop codon due to conserved overlapping splice site
<sup>b</sup> Unable to locate a stop codon. The structure of the final two exons is conserved among human, mouse and rat with the ultimate residue overlapping the splice site. The final coding exon therefore only contains two bp and the stop codon making it hard to determine the location of the final exon in the gerbil without supporting RNA information.
<sup>c</sup> Missing 99 residues (121-219) due to assembly gap
<sup>d</sup> Start location is that of exon 2 as exon 1 is missing due to an assembly gap.
<sup>e</sup> The cytoplasmic tail of $\beta 2$ genes are encoded by an exon with no known homology to other exons in the MHC II genes and has a low degree of homology between mouse and rat [42]. There is therefore some uncertainty related to the completeness of the final exon of *Rhop-DRB2*.
Legend: The table details the assigned great gerbil gene names, which scaffold and in what orientation they are located as well as genomic location on the scaffold for start and end of the genes. Information on the size of the translated amino acid sequence and whether it is complete is also shown.

Any similar duplication of the Rhop-DRB1 gene is not seen in either of the two close family members of the Gerbillinae subfamily used in our comparative analyses. BLAST searches of the sand rat genome returned a single full-length copy of the *β*1 gene and a near full-length copy of the *β*2 gene (Fig. 5 and Additional file 2: Table S6). According to the annotations of the Mongolian gerbil genome provided by NCBI, this species contains two copies of the DR locus *β* genes. A manual tBLASTn search using the protein sequences of Mongolian gerbil DRB genes to search the genome assembly did not yield additional hits of *β* genes in this locus that could have been missed in the automatic annotation process. The phylogeny confirms the copies found in Mongolian gerbil to be *β*1 and *β*2 genes (Fig. 5).

### *MHCII DRB* promoters

*MHCII* genes each contain a proximal promoter with conserved elements (S-X-Y motifs) that are crucial for the efficient expression of the gene. We aligned the proximal promoter of the *β* genes of the DR locus in great gerbil and the other investigated species to establish if the integrity of the promoter was conserved as well as examining similarities and potential dissimilarities causing the previously reported differences in transcription and expression of *β*1 and *β*2 genes in rodents [42,44]. The alignment of the promoter region reveals the conserved structure and similarities within *β*1 and *β*2 genes as well as characteristic differences (Fig. 6 and Table 4). Clear similarities are seen for the proximal promoter regions of *Rhop-DRB1* and *Rhop-DRB3* to the other rodent and human *β*1 promoters, as illustrated by high sequence similarity and the presence of a CCAAT box just downstream of the Y motif in all investigated rodent *β*1 promoters. Notably, the CCAAT box is missing in *β*2 promoters. The crucial distance between the S and X motifs is conserved in all *β* genes and the integrity of the S-X-Y motifs is observable for *Rhop-DRB1* and *DRB3* promoters. However, both the S and X box of *DRB2* are compromised by deletions in great gerbil. The deletion in the X box severely disrupt the motif and reduce its size by half. An identical deletion in the X box is seen in Mongolian gerbil while the sand rat X box sequence covering the deleted parts is highly divergent from the conserved sequence found in the rest of the promoters (Fig. 6). Furthermore, for the *β*2 genes, two deletions downstream of the motifs are shared among all rodents in the alignment as well as an additional insertion observed in Gerbillinae members.

**FIG. 6.**
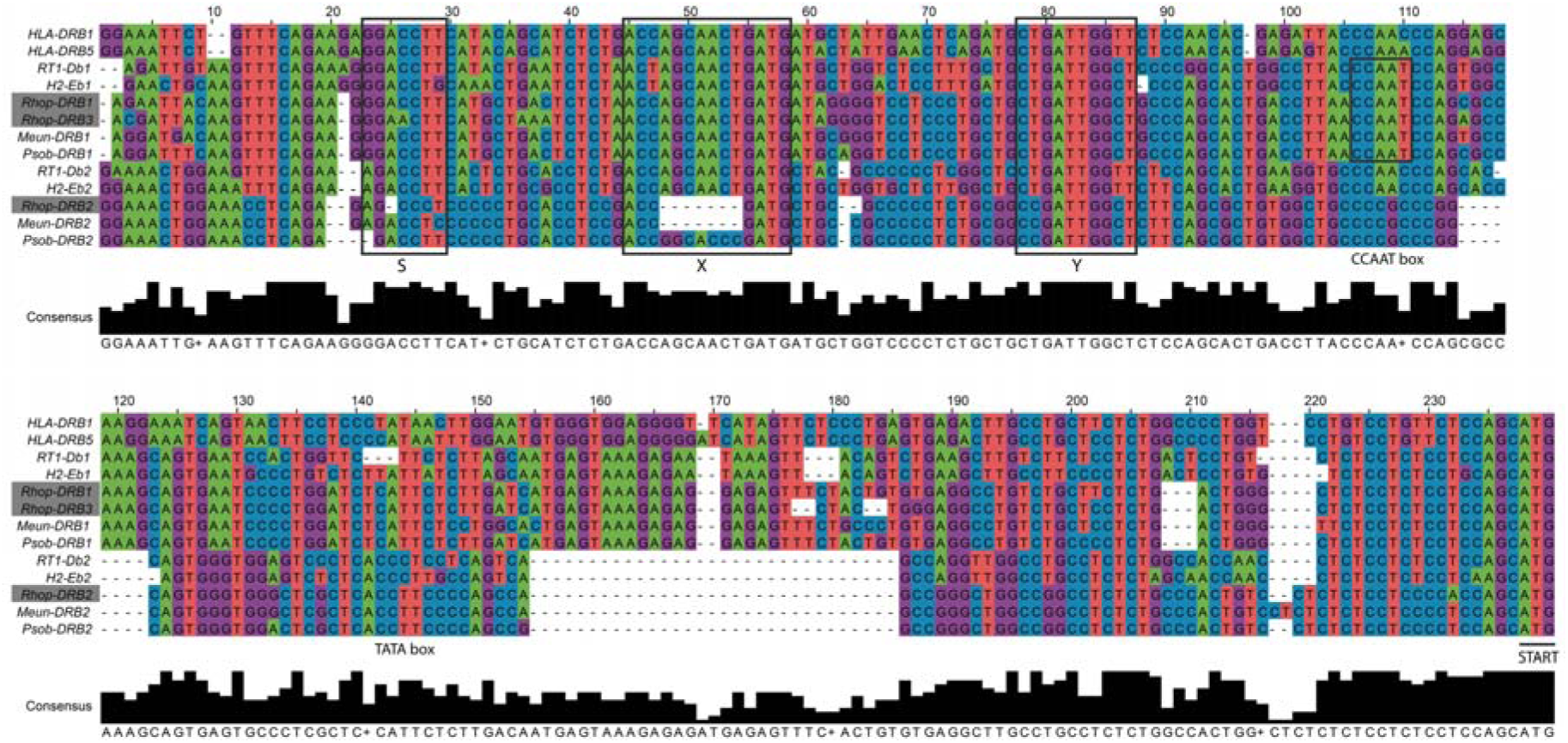
Alignment of the DR locus β1 and β2 proximal promoters. Sequences of the proximal promoters of *β*1 and *β*2 genes of the DR locus (E locus in mouse and D locus in rat) were aligned in MEGA7 [45] using MUSCLE with default parameters. The resulting alignment was edited manually for obvious misalignments and transferred and displayed in Jalview [46]. For visualization purposes only, the alignment was further edited in Adobe Illustrator (CS6), changing colors of the bases and adding boxes to point out the S-X-Y motifs. The three copies of DRB genes located in the great gerbil genome are marked with grey boxes. The alignment shows clear similarities of the proximal promoter region of *Rhop-DRB1* and *Rhop-DRB3* to the other rodent and human *β*1 promoter sequences. For the *DRB2* genes, two deletions are shared among all rodents in the alignment as well as additional indels observed in Gerbillinae members. Most notably, both great gerbil and Mongolian gerbil have deletions of half the X box while sand rat X box sequences in that same position is highly divergent from the otherwise conserved sequence seen in the alignment.

**Table 4.** Coordinates of *MHCII* S-X-Y motifs of the promoters of the DRB genes in investigated species.

| Gene | S-X spacing (bp) | X-Y spacing (bp) | S-X-Y Relative position (S-X-Y end to gene start) | CCAAT box |
| --- | --- | --- | --- | --- |
| *HLA-DRB1* | 15 | 19 | -145 | TATA box |
| *HLA-DRB5* | 15 | 19 | -146 |  |
| *RT1-Db1* | 15 | 19 | -137 | Yes |
| *H2-Eb1* | 15 | 19 | -139 | Yes + TATA box |
| *Rhop-DRB1* | 15 | 19 | -141 | Yes |
| *Rhop-DRB3* | 15 | 19 | -137 | Yes |
| *Meun-DRB1* | 15 | 19 | -141 | Yes |
| *Psob-DRB1* | 15 | 19 | -141 | Yes |
| *RT1-Db2* | 15 | 18 | -110 |  |
| *H2-Eb2* | 15 | 19 | -110 |  |
| *Rhop-DRB2* | 15 | 17 | -109 |  |
| *Meun-DRB2* | 15 | 18 | -111 |  |
| *Psob-DRB2* | 15 | 18 | -109 |  |

Legend: The overview details the relative coordinates for conserved motifs in the promoter regions of great gerbil (*Rhop-DRB*), sand rat (*Psob-DRB*), Mongolian gerbil (*Meun-DRB*), human (*HLA-DRB*), mouse (*H2-Eb*) and rat (*RT1-Db*) orthologous genes. The S-X and X-Y spacing refers to the distance between the S and X, and X and Y motifs, respectively. The S-X-Y Relative position show the distance (bp) between the end of the Y motif and the start codon of the gene.

### Peptide binding affinity predictions and expression of Rhop-DR MHCII molecules

Mouse and rat β2 molecules have been shown to have an extraordinary capacity to present the *Y. pseudotuberculosis* superantigen mitogen (YPm) [42]. We therefore investigated the peptide binding affinities of the Rhop-DR molecules by running translations of *Rhop-DRA* in combinations with each of the three *Rhop-DRB* genes through the NetMHCIIpan 3.2 server [47] along with peptide/protein sequences of YPm, *Y. pestis* F1 ‘capsular’ antigen and LcrV antigen. Universally, the Rhop-DRB3 shows an affinity profile identical to that of Rhop-DRB2 displaying high affinity towards both *Y. pseudotuberculosis* and *Y. pestis* epitopes while Rhop-DRB1 does not (Fig.7 and Additional file 3). The translated great gerbil MHCII from DP and DQ loci were also tested for peptide binding affinity but only Rhop-DP displayed affinity to one of the epitopes tested. Furthermore, analyses of the translated amino acid sequences of sand rat DR (Psob-DR) molecules as well as published protein sequences of Mongolian gerbil DR (Meun-DR) molecules and the mouse ortholog H2-E confirmed the high affinity of β2 molecules to *Y. pseudotuberculosis* and *Y. pestis* (Additional file 1: Figure S10 and Additional file 3). The equal capacity of Rhop-DRB2 and Rhop-DRB3 to putatively present *Yersiniae* combined with the proximal promoter investigations lead us to question the expression of DRB genes in great gerbil. Searching a set of raw counts of great gerbil expressed genes, reveal that Rhop-DRB1 and Rhop-DRB3 are both similarly expressed at similar levels while Rhop-DRB2 is not expressed or at undetectable levels (3936 and 2279 vs 14).

**FIG. 7.**
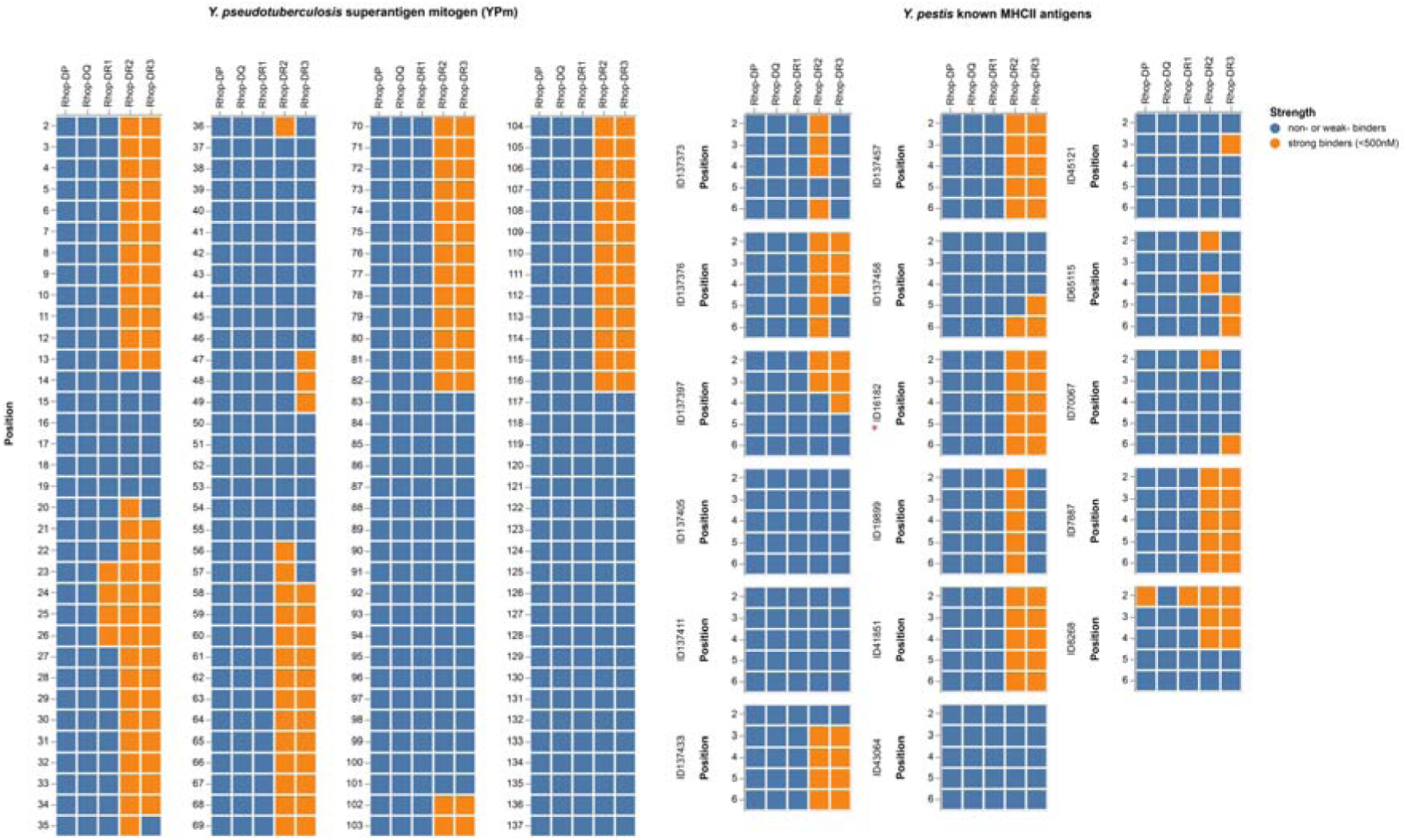
Affinity predictions of great gerbil MHCII molecules. Affinity predictions of great gerbil MHC class II molecules represented as a heatmap. For the known *Y. pestis* antigens all are from the F1 capsule precursor except ID16182 (red asterix) which is from the V antigen. Strong binders are defined as <500 nM and depicted in orange, while weak or non-binders are represented in blue.

## Discussion

Here we present a highly contiguous *de novo* genome assembly of the great gerbil covering over 96 % of the estimated genome size and almost 88 % of the gene space, which is equivalent to the genic completeness reported in the recently published and close relative sand rat genome [48] (Additional file 2: Table S7). By comparative genomic analyses where we include genome data from its close relatives within the Gerbillinae, we provide novel insight into the innate and adaptive immunological genomic landscape of this key plague host species.

### The *TLR* repertoire in the great gerbil and Gerbillinae

*TLRs* are essential components of PRRs and the innate immune system as they alert the adaptive immune system of the presence of invading pathogens [49]. The detailed characterization of *TLRs* did not uncover any species-specific features for the great gerbil. However, a shared *TLR* gene repertoire for the Gerbillinae lineage (i.e. the great gerbil, sand rat and Mongolian gerbil), with gene losses of *TLR8*, *TLR10* and all members of the *TLR11*-subfamily was revealed. This finding could indicate quite similar selective pressures on these species, at least in regard to their function of *TLRs*, all being desert dwelling, burrowing rodents living in arid or semi-arid ecosystems and being capable of carrying plague. Thus, it is possible that the members of this clade have reduced the *TLR* repertoire in a cost-benefit response to environmental constraints or due to altered repertoire of pathogen exposure [50]. These results are in line with the fairly conserved *TLR* gene repertoire reported within the vertebrate lineage [51], although the repertoire of *TLR* genes present within vertebrate groups can show major differences [51-53], presumably in response to presence or lack of certain pathogen or environmental pressures [54,55]. Outside of Gerbillinae, the presence of TLR11-subfamily appears to be universal in Rodentia, however functionally lost from the human repertoire [51,53]. The TLR11-subfamily recognizes parasites and bacteria through profilin, flagellin and 23S ribosomal RNA [56,57] and it is possible that cross-recognition of these patterns by other TLR members or other PRRs might have made the TLR11-subfamily redundant in Gerbillinae and humans [50]. The varying degree of point mutations, frameshift-causing indels and in some cases almost complete elimination of sequence in *TLR8*, *TLR10* and *TLR11-13* in Gerbillinae suggest successive losses of these receptors, where a shared pseudogenization of *TLR12-13* across all species investigated were recorded. For *TLR11* however, the pseudogenization seems to have occurred in multiple steps, i.e. with a more recent event in the great gerbil where a near full-length sequence was identified compared to the shorter fragments identified for Mongolian gerbil and sand rat (Additional file 1: Figure S2C). Furthermore, the high degree of shared disruptive mutations among all three species of Gerbillinae indicates that the initiation of pseudogenization predates the speciation estimated to have occurred about 5.5 Mya [58].

In the context of plague susceptibility, the branch specific diversifying selection reported here for *TLR7* and *TLR9* in Gerbillinae is intriguing, as both receptors have been implicated to affect the outcome of plague infection in mice and humans [59-61]. For instance, the study by Dhariwala et al. (2017) showed, in a murine model, that TLR7 recognizes intracellular *Y. pestis* and is important for defense against disease in the lungs but was detrimental to septicemic plague [59]. Moreover, recognition of *Y. pestis* by TLR9 was also demonstrated by Saikh et al. (2009) in human monocytes [61]. All but one of the residues under site specific selection seen in *TLR7* and *TLR9* were located in the ectodomain, which may suggest possible alterations in ligand recognition driven by selection pressure from *Y. pestis* or other shared pathogens. Stimulation of *TLR7* and *TLR9* have also been reported to regulate antigen presentation by *MHCII* in murine macrophages [62]. These data could therefore indicate a possible connection of the selection in *TLR7* and *TLR9* with the great gerbil duplication in *MHCII*. For *TLR4*, the selection tests and sequence alignment analysis did not reveal any branch-specific selection for great gerbil nor Gerbillinae, whereas we did detect signs of site-specific selection in the ectodomain that occasionally was driven by great gerbil or Gerbillinae substitutions. TLR4 is the prototypical PRR for detection of lipopolysaccharides (LPS) found in the outer membrane of gram-negative bacteria like *Y. pestis*. As part of the arms race, however, it is well known that gram-negative bacteria, including *Y. pestis*, alter the conformation of their LPS in order to avoid recognition and strong stimulation of the TLR4-MD2-CD14 receptor complex [63-65]. Despite this, in mice at least, some inflammatory signaling still occurs through this receptor complex but require particular residues in TLR4 not found to be conserved in the Gerbillinae lineage. Whether other mutations in TLR4 in Gerbillinae have a similar functionality as the residues that allow mice to respond to *Y. pestis* LPS is not known. However, if such functionality is missing in Gerbillinae, the loss of responsiveness to the hypoacetylated LPS [19] could perhaps defer some protection from pathologies caused by excessive initiation of inflammatory responses [66], and thus TLR4 is not likely directly involved in the resistance of plague in great gerbils.

Cumulatively, our investigations of the great gerbil innate immune system, focusing on the *TLR* gene repertoire, reveal shared gene losses within *TLR* gene families for the Gerbillinae lineage, all being desert dwelling species capable of carrying plague. The evolutionary analyses conducted did not uncover any great gerbil-specific features that could explain their resistance to *Y. pestis*, indicating that other PRRs (not investigated here) could be more directly involved during the innate immune response to plague infection in the great gerbil [36].

### Great gerbil MHC repertoires

MHCI and II proteins are crucial links between the innate and adaptive immune system continuously presenting peptides on the cell surface for recognition by CD8+ and CD4+ T cells respectively, and MHC genes readily undergo duplications, deletions and pseudogenization [67]. For *MHCI*, the discovery of 16 copies in great gerbil is in somewhat agreement with what has earlier been reported in rodents, where the MHCI region is found to have undergone extensive duplication followed by sub-and neofunctionalization with several genes involved in non-immune functions [68,69]. However, it should be noted that our copy number estimation is most likely an underestimation, due to the assembly collapse in almost all *MHCI* containing regions identified. Furthermore, not all copies could be confidently placed in the gene maps as some scaffolds lacked colocalizing framework genes. These two factors are the probable reason why the great gerbil appears to be lacking some *MHCI* genes compared to mouse and rat.

For *MHCII* we discovered a gerbil-specific duplication that is not present in other closely related plague hosts or in other rodents investigated. The phylogeny established the duplication’s (*Rhop-DRB3*) relationship to *Rhop-DRB1* and other mammalian *β*1 genes and reflects the orthology of mammalian *MHCII* genes [70]. The localization of *Rhop-DRB3* outside of the generally conserved framework of the *MHCII* region and not in tandem with the other *β* genes of the *DR* locus is unusual and is not generally seen for eutherian mammals. For instance, major duplication events with altered organization and orientation of *DR* and *DQ* genes has been reported for the *MHCII* region in horse (*Equus caballus*), however all genes are found within the framework genes [71]. Duplications tend to disperse in the genome as they age [72], thus the reversed orientation and translocation of the great gerbil copy might indicate that the duplication event is ancient occurring sometime after the species split approximately 5 Mya. However, it must also be noted that there are several assembly gaps located between *Rhop-DRB1* and *Rhop-DRB3* resulting in the possibility of the translocation being a result of an assembly error.

Predictions of the affinity of the β1, β2 and β3 MHCII molecules to *Y. pestis* and *Y. pseudotuberculosis* antigens matched the reported high affinity of rodent β2 molecules for *Yersiniae epitopes* [42]. Rhop-DRB3 had an equally high affinity and largely identical affinity-profile as Rhop-DRB2. A high affinity for *Y. pestis* epitopes is important in the immune response against plague, as the initiation of a T cell response is more efficient and requires fewer APCs and T cells when high-affinity peptides are presented by MHCII molecules [73]. In the early stages of an infection where presence of antigen is low, there will be fewer MHCII molecules presenting peptides and affinity for those peptides is paramount to fast initiation of the immune response against the pathogen. Individuals presenting MHCII molecules with high affinity for pathogen epitopes are able to raise an immune defense more quickly and have a better chance of fighting off the rapidly progressing infection than individuals that are fractionally slower. This fractional advantage could mean the difference between death or survival.

We find comparable expression levels for *Rhop-DRB1* and *Rhop-DRB3* but no detectable expression of *Rhop-DRB2*. These similarities and differences are likely explained by the variations discovered in the proximal promoter of the genes. Integrity of the conserved motifs and the spacing between them is necessary for assembly of the enhanceosome complex of transcription factors and subsequent binding of Class II Major Histocompatibilty Complex transactivator (*CIITA*), and is essential for efficient expression of *MHCII* genes. The conservation of the proximal promoter of *Rhop-DRB3* along with the overall sequence similarity with other *β*1 genes are indicative of a similar expression pattern. In contrast, the deletion in the X box of *Rhop-DRB2* reducing the motif to half the size will likely affect the ability of the transcription factors to bind and could explain the lack of expression. Similar disruptions in the *β*2 genes of the other Gerbillinae were found along with a major deletion further downstream in all *β*2 genes that perhaps explains the previously reported low and unusual pattern of transcription for rodent *β*2 genes [42,44]. The equal affinity profile but different expression levels of *Rhop-DRB2* and *Rhop-DRB3* could mean that *Rhop-DRB3* has taken over the immune function lost by the lack of expression of *Rhop-DRB2*. The selective pressure might have come from *Yersinia* or pathogens similar to *Yersiniae*. A nonclassical function of MHCII molecules have also been reported where intracellular MHCII interacted with components of the *TLR* signaling pathway in a way that suggested MHCII molecules are required for full activation of the TLR-triggered innate immune response [74]. Moreover, in vertebrates the *MHCII* DRB genes are identified as highly polymorphic and specific allele variants have frequently been linked to increased susceptibility to diseases in humans [75]. Intriguingly, in a recent study by Cobble et al. (2016) it was suggested that allelic variation of the *DRB1* locus could be linked to plague survival in Gunnison’s prairie dog colonies [40]. Thus, investigating how the genetic variation of the *DRB1* and *DRB3* loci in great gerbil manifests at the population level and the affinity of these allelic variants to *Yersiniae* epitopes, would be the next step to further our understanding of the plague resistant key host species in Central Asia.

From the analyses conducted on the genomic landscape of the adaptive immune system of the great gerbil, i.e. *MHCI* and *MHCII* more specifically, the most interesting reporting is the duplication of an *MHCII* gene. *In silico* analyses of *Rhop-DRB3* indicate a high predicted affinity for *Y. pestis* epitopes, which may result in faster initiation of the adaptive immune system in great gerbils when exposed to the pathogen, and thus could explain the high degree of plague resistance in this species.

## Conclusion

Plague has historically had a vast impact on human society through major pandemics, however it mainly circulates in rodent communities. A key issue is to understand host-pathogen interactions in these rodent hosts. From the pathogen-perspective, research has studied how *Y. pestis* has evolved to evade both detection and destruction by the mammalian immune system to establish infection. In this study, we have demonstrated the power of using whole genome sequencing of a wild plague reservoir species to gain new insight into the genomic landscape of its resistance by immuno-comparative analyses with closely related plague hosts and other mammals. We reveal the duplication of an MHCII gene in great gerbils with a computed peptide binding profile that putatively would cause a faster initiation of the adaptive immune system when exposed to *Yersiniae* epitopes. We also find signs of positive selection in *TLR7* and *TLR9*, which have been shown to regulate antigen presentation and impact the outcome of a plague infection. Investigations into how the genetic variation of the MHCII locus manifests at the population level are necessary to further understand the role of the gene duplication in the resistance of plague in great gerbils. Comprehending the genetic basis for plague resistance is crucial to understand the persistence of plague in large regions of the world and the great gerbil *de novo* genome assembly is a valuable anchor for such work, as well as a resource for future comparative work in host-pathogen interactions, evolution (of resistance) and adaptation.

## Methods

### Sampling and sequencing

A male great gerbil weighing 180g was captured in the Midong District outside Urumqi in Xinjiang Province, China in October 2013. The animal was humanely euthanized and tissue samples of liver were conserved in ethanol prior to DNA extraction. Blood samples from the individual were screened for F1 ‘capsular’ antigen (Caf1) and anti-F1 as described in [30,76] to confirm plague negative status. The DNA used in the library construction was extracted from liver tissue using Gentra Puregene Tissue Kit (Qiagen Inc. USA). Use of great gerbil tissue was approved by the Committee for Animal Welfares of Xinjiang CDC, China.

The sequence strategy was tailored towards the ALLPATHS-LG assembly software (Broad Institute, Cambridge, MA) following their recommendations for platform choice and fragment size resulting in the combination of one short paired-end (PE) library with an average insert size of 220 bp (150 bp read length) and two mate-pair (MP) libraries of 3 kbp and 10 kbp insert size (100 bp read length). See Additional file 2: Table S1 for a list of libraries and sequence yields. Sequencing for the *de novo* assembly of the great gerbil reference genome was performed on the Illumina platform using HiSeq2500 instruments at the Norwegian Sequencing Centre at the University of Oslo for the PE library (https://www.sequencing.uio.no) and using HiSeq2000 instruments at Génome Québec at McGill University for the MP libraries (http://gqinnovationcenter.com/index.aspx?l=e).

### Genome assembly and Maker annotation

The Illumina sequences were quality checked using FastQC v0.11.2 and SGA-preqc (downloaded 25^th^ June 2014) with default parameters. Both MP libraries were trimmed for adapter sequences using cutadapt v1.5 with option-b and a list of adapters used in MP library prep [77] and the trimmed reads were used alongside the PE short read as input for ALLPATHS-LG v48639 generating a *de novo* assembly. This combination of short-read sequencing technology combined with the ALLPATHS-LG assembly algorithm is documented to perform well in birds and mammals [78-80]. File preparations were conducted according to manufacturer’s recommendation and the option TARGETS=submission was added to the run to obtain a submission prepared assembly version.

Assembly completeness was assessed by analysing the extent of conserved eukaryotic genes present using CEGMA v2.4.010312 and BUSCO v1.1.b [81-83]. Gene mining for the highly conserved Homeobox (*HOX*) genes was also conducted as an additional assessment of assembly completeness (see Additional file 1: Note S1 and Figure S10). All reads were mapped back to the assembly using BWA-MEM v 0.7.5a and the resulting bam files were used alongside the assembly in REAPR v 1.0.17 to evaluate potential scaffolding errors as well as in Blobology to inspect the assembly for possible contaminants, creating Taxon-Annotated-GC-Coverage (TAGC) plots of the results from BLAST searches of the NCBI database [84].

The genome assembly was annotated using the MAKER2 pipeline v2.31 run iteratively in a two-pass fashion (as described in https://github.com/sujaikumar/assemblage/blob/master/README-annotation.md) [85]. Multiple steps are required prior to the first pass though MAKER2 and include creating a repeat library for repeat masking and training three different ab initio gene predictors. Firstly, construction of the repeat library was conducted as described in [86]. In brief, a *de novo* repeat library was created for the assembly by running RepeatModeler v1.0.8 with default parameters, and sequences matching known proteins of repetitive nature were removed from the repeat library through BLASTx against the UniProt database. Next, GeneMark-ES v2.3e was trained on the genome assembly using default parameters with the exception of reducing the–min-contig parameter to 10.000 [87]. SNAP v20131129 and AUGUSTUS v3.0.2 was trained on the genes found by CEGMA and BUSCO, respectively. The generated gene predictors and the repeat library were used in the first pass alongside proteins from UniProt/SwissProt (downloaded 16^th^ February 2016) as protein homology evidence and Mus musculus cDNA as alternative EST evidence (GRCm38 downloaded from Ensembl). For the second pass, SNAP and AUGUSTUS were retrained with the generated MAKER2 predictions and otherwise performed with the same setup. The resulting gene predictions had domain annotations and putative functions added using InterProScan v5.4.47 and BLASTp against the UniProt database with evalue 1e-5 (same methodology as [86,88]). Finally, the output was filtered using the MAKER2 default filtering approach only retaining predictions with AED <1.

### Genome mining and gene alignments

We searched for *TLR* genes, associated receptors and adaptor molecules as well as genes of the MHC region (complete list of genes can be found in Additional file 2: Table S4) collected from UniProt and Ensembl. Throughout, we performed tBLASTn searches, manual assembly exon by exon in MEGA7 and verified annotations through reciprocal BLASTx against the NCBI database and phylogenetic analysis including orthologues from human (*Homo sapiens*), mouse (*Mus musculus*), rat (*Rattus norvegicus*) and all three members of the *Gerbillinae* subfamily. For details on the phylogenetic analyses we refer to descriptions in sections below. In the *TLR* analyses Algerian mouse (*M. spretus*), Ryukyu mouse (*M. caroli*), Chinese hamster (*Cricetulus griseus*) and Chinese tree shrew (*Tupaia belangeri chinensis*) were also included.

Sand rat and Mongolian gerbil genome assemblies were downloaded from NCBI (September 12^th^ 2017). The genome assemblies of the great gerbil, sand rat and Mongolian gerbil were made into searchable databases for gene mining using the makeblastdb command of the blast+ v2.6.0 program. Local tBLASTn searches, using protein sequences of mouse and occasionally rat, human and Mongolian gerbil as queries, were executed with default parameters including an e-value cut-off of 1e+1. The low e-value was utilized to capture more divergent sequence homologs. Hits were extracted from assemblies using bedtools v2.26.0 and aligned with orthologs in MEGA v7.0.26 using MUSCLE with default parameters. In cases where annotations for some of the *TLRs* for a species were missing in Ensembl and could not be located in either the NCBI nucleotide database or in UniProt, the Ensembl BLAST Tool (tBLASTn) was used with default parameters to find the genomic region of interest using queries from mouse.

### Synteny analyses of MHC regions

A combination of the Ensembl genome browser v92 and comparisons presented in [89] and tBLASTn searches, as described above, were used in synteny analyses of the MHCI and II regions of human, rat and mouse with great gerbil. Synteny of MHCII genes of sand rat and Mongolian gerbil were also investigated, however for simplicity and visualization purposes not included in the figure (Fig. 3).

### Alignment and phylogenetic reconstruction of TLR and MHC

Sequences were aligned with MAFFT [90] using default parameters: for both nucleotides and amino acid alignments the E-INS-i model was utilized. The resulting alignments were edited manually using Mesquite v3.4 [91]. See Additional file 2: Tables S8-10 for accession numbers.

Ambiguously aligned characters were removed from each alignment using Gblocks [92] with the least stringent parameters for codons and proteins.

Maximum likelihood (ML) phylogenetic analyses were performed using the “AUTO” parameter in RAxML v8.0.26 [93] to establish the evolutionary model with the best fit. The general time reversible (GTR) model was the preferred model for the nucleotide alignments, and JTT for the amino acid alignments. The topology with the highest likelihood score of 100 heuristic searches was chosen. Bootstrap values were calculated from 500 pseudoreplicates. Taxa with unstable phylogenetic affinities were pre-filtered using RogueNaRok [94] based on evaluation of a 50 % majority rule (MR) consensus tree, in addition to exclusion of taxa with >50 % gaps in the alignment.

Bayesian inference (BI) was performed using a modified version of MrBayes v3.2 [95] (https://github.com/astanabe/mrbayes5d). The dataset was executed under a separate gamma distribution. Two independent runs, each with three heated and one cold Markov Chain Monte Carlo (MCMC) chain, were started from a random starting tree. The MCMC chains were run for 20,000,000 generations with trees sampled every 1,000^th^ generation. The posterior probabilities and mean marginal likelihood values of the trees were calculated after the burn-in phase (25 %), which was determined from the marginal likelihood scores of the initially sampled trees. The average split frequencies of the two runs were < 0.01, indicating the convergence of the MCMC chains.

### Selection analyses

All full-length *TLRs* located in the genomes of great gerbil, sand rat and Mongolian gerbil along with other mammalian *TLRs* (Additional file 2: Table S8) were analysed in both classic Datamonkey and Datamonkey 2.0 (datamonkey.org) testing for signs of selection with a phylogeny guided approach [96,97]. For each *TLR* gene alignment a model test was first run prior to the selection test and the proposed best model was used in the analyses. The mixed effects model of evolution (MEME) and adaptive branch-site random effects model (aBSREL) were used to test for site based and branch level episodic selection, respectively [98-100]. aBSREL was iterated three times per gene alignment, initially running an exploratory analysis were all branches were tested for positive selection and subsequently in a hypothesis mode by which first the Gerbillinae clade and secondly the great gerbil was selected as “foreground” branches to test for positive selection. All *TLR* alignments are available in the Github repository (https://github.com/uio-cels/Nilsson_innate_and_adaptive).

### TLR protein structure prediction

Translated full-length great gerbil *TLR* sequences were submitted to the Phyre2 structure prediction server for modelling [101]. All sequences were modelled against human TLR5 (c3j0aA) and the resulting structures were colored for visualization purposes using Jmol (Jmol: an open-source Java viewer for chemical structures in 3D. http://www.jmol.org/). Colors were used to differentiate between helices, sheets and loops as well as the transmembrane domain, linker and TIR domain. Sites found in the MEME selection analysis were indicated in pink and further highlighted with arrows (Additional file 1: Figures S7-9). All great gerbil PDB files are available in the GitHub repository (https://github.com/uio-cels/Nilsson_innate_and_adaptive).

As TLR4 is the prototypical PRR for lipopolysaccharide (LPS) which are found in all gram-negative bacteria including *Y. pestis*, we subjected the sequence alignment to additional investigation of certain residues indicated in the literature to have an impact on signaling [19]. These were the residues at position 367 and 434, which in mouse are both basic and positively charged, enabling the mouse TLR4 to maintain some signaling even for hypoacetylated LPS [19]. Hypoacetylated LPS is a common strategy for gram-negative bacteria to avoid recognition and strong stimulation of the TLR4-MD2-CD14 receptor complex [63-65].

### MHCII promoter investigation

The region 400 bp upstream of human HLA-DRB, mouse H2-Eb and rat RT-Db genes were retrieved from Ensembl (GRch38.p12, GRCm38.p6 and Rnor_6.0). Similarly, the region 400 bp upstream of the start codon of DRB genes in the three Gerbillinae were retrieved using bedtools v2.26.0. Putative promoter S-X-Y motifs, as presented for mouse in [102], were manually identified for each gene in MEGA7 and all sequences were subsequently aligned using MUSCLE with default parameters [102].

### Peptide binding affinity

The functionality of MHCII genes is defined by the degree of expression of the MHC genes themselves, and the proteins ability to bind disease-specific peptides to present to the immune system. The ability of an MHCII protein to bind particular peptides can with some degree of confidence be estimated by MHC prediction algorithms, even for unknown MHCII molecules, as long as the alpha and beta-chain protein sequences are available [47]. We here use the NetMHCIIpan predictor v3.2 [47] to estimate the peptide binding affinities of the novel Rhop-DRB3 MHCII molecule and compare it to various other MHCII molecules from great gerbil, mouse, sand rat and Mongolian gerbil. The program was run with default settings and provided with the relevant protein sequences of alpha and beta chains. We compared the predicted binding affinity of these MHCII molecules for 17 known *Y. pestis* epitopes derived from positive ligand assays of *Y. pestis* (https://www.iedb.org/). Specifically, we tested against 16 ligands derived from the F1 capsule antigen of *Y. pestis*, and 1 ligand from the virulence-associated Low calcium V antigen (LcrV) of *Y. pestis*. In addition, we compared the binding affinity of these MHCII molecules against the superantigen *Y. pseudotuberculosis* derived mitogen precursor (YPm) [42]. The threshold for binders was set to <500nM [47].

### RNA sampling and sequencing

Two additional great gerbils were captured in the Midong District outside Urumqi in Xinjiang Province, China, in September 2014. The animals were held in captivity for 35 days before being humanely euthanized and liver tissue samples were conserved in RNAlater^TM^ at −20 °C prior to RNA extraction. RNA was extracted using standard chloroform procedure [103]. Library prep and sequencing were conducted at the Beijing Genomics Institute (BGI, https://www.bgi.com/us/sequencing-services/dna-sequencing/) using Illumina TruSeq RNA Sample Prep Kit and PE sequencing on the HiSeq4000 instrument (150 bp read length).

The reads were trimmed using trimmomatic v0.36 and mapped to the genome assembly using hisat2 v2.0.5 with default parameters. A raw count matrix was created by using htseq v0.7.2 with default parameters to extract the raw counts from the mapped files.

## Acknowledgements

All computational work was performed on the Abel Supercomputing Cluster (Norwegian metacenter for High Performance Computing (NOTUR) and the University of Oslo) operated by the Research Computing Services group at USIT, the University of Oslo IT-department and the Cod nodes of CEES. Sequencing library creation and high throughput sequencing was carried out at the Norwegian Sequencing Centre (NSC), University of Oslo, Norway, and McGill University and Genome Quebec Innovation Centre, Canada.

We would like to thank Morten Skage for assistance in sequence library construction and Ole K. Tørresen, Srinidhi Varadharajan, Tore O. Elgvin and Cassandra N. Trier for helpful advice and support during assembly and annotations steps of the genome, Helle T. Baalsrud for advice during genome mining and Tone F. Gregers for helpful discussions regarding MHCII. For early access to the sand rat genome assembly we thank John F. Mulley.

## Availability of data and materials

The genome assembly has been deposited at DDBJ/ENA/GenBank under the accession REGO00000000. The version described in this paper is version REGO01000000.

The genome assembly and annotation are also available from FigShare: In the following GitHub repository are files of immune gene alignments, PDB files and more: https://github.com/uio-cels/Nilsson_innate_and_adaptive

## Authors’ contributions

PN created the genome assembly and annotated it, performed all BLAST-based, *TLR* based and promoter analysis and wrote the first draft of the manuscript. MHS conducted the protein model analyses of *TLRs* and assisted in the BLAST-based and *TLR* analyses. BVS performed the MHCII affinity analyses. RJSO performed phylogenetic analysis of TLR, MHCI and MHCII genes. YZ, sampled, acclimatised and tested individual great gerbil for plague. RL, YC and YS extracted DNA and RNA for sequencing. PN, WRE, BVS, SJ and KSJ designed the sequencing strategy. WRE, BVS, SJ, KSJ, NCS and RY oversaw the project. All authors read and approved the final manuscript.

## Ethics approval

Use of great gerbil tissue was approved by the Committee for Animal Welfares of Xinjiang Centre for Disease Control and Prevention, China. Sampling was performed prior to Chinas signature of the Nagoya Protocol (date of accession September 6^th^ 2016). The sampled species have a “least concern” status in the IUCN Red List of Threatened Species.

## Consent for publication

Not applicable.

## Additional files

Additional file 1: Additional figures and one Note detailing the HOX gene mining (DOCX 19.2Mb)

Additional file 2: Additional tables (DOCX 66Kb)

Additional file 3: Peptide binding affinity predictions for all MHCII molecules run in NetMHCIIpan predictor v3.2 (XLSX)

## References

1. Anisimov AP, Lindler LE, Pier GB. Intraspecific Diversity of Yersinia pestis. Clinical Microbiology Reviews. 2004;17:434–64.

2. Addink EA, De Jong SM, Davis SA, Dubyanskiy V, Burdelov LA, Leirs H. The use of high-resolution remote sensing for plague surveillance in Kazakhstan. Remote Sensing of Environment. Elsevier; 2010;114:674–81.

3. Nowak RM. Walker’s Mammals of the World. JHU Press; 1999.

4. Zhang Z, Zhong W, Fan N. Rodent problems and management in the grasslands of China. In: Singleton GR, Hinds LA, Krebs CJ, Spratt DM, editors. Rats, mice and people: rodent biology and management. researchgate.net; 2003. pp. 316–9.

5. Gage KL, Kosoy MY. Natural history of plague: perspectives from more than a century of research. Annu. Rev. Entomol. 2005;50:505–28.

6. Yang R, Anisimov A. Yersinia pestis: Retrospective and Perspective. Springer; 2016.

7. Stenseth NC, Atshabar BB, Begon M, Belmain SR, Bertherat E, Carniel E, et al. Plague: past, present, and future. PLoS Med. Public Library of Science; 2008;5:e3.

8. Bramanti B, Stenseth NC, Walløe L, Lei X. Plague: A Disease Which Changed the Path of Human Civilization. In: Yang R, Anisimov A, editors. Yersinia pestis: Retrospective and Perspective. Dordrecht: Springer; 2016. pp. 1–26.

9. Hinnebusch BJ, Jarrett CO, Bland DM. “Fleaing” the Plague: Adaptations of Yersinia pestis to Its Insect Vector That Lead to Transmission. Annu. Rev. Microbiol. Annual Reviews; 2017;71:215–32.

10. Samia NI, Kausrud KL, Heesterbeek H, Ageyev V, Begon M, Chan K-S, et al. Dynamics of the plague–wildlife–human system in Central Asia are controlled by two epidemiological thresholds. Proceedings of the National Academy of Sciences. National Academy of Sciences; 2011;108:14527–32.

11. Nguyen VK, Parra-Rojas C, Hernandez-Vargas EA. The 2017 plague outbreak in Madagascar: Data descriptions and epidemic modelling. Epidemics. 2018.

12. Boisier P, Rahalison L, Rasolomaharo M, Ratsitorahina M, Mahafaly M, Razafimahefa M, et al. Epidemiologic Features of Four Successive Annual Outbreaks of Bubonic Plague in Mahajanga, Madagascar. Emerging Infect. Dis. Centers for Disease Control and Prevention; 2002;8:311–6.

13. Migliani R, Chanteau S, Rahalison L, Ratsitorahina M, Boutin JP, Ratsifasoamanana L, et al. Epidemiological trends for human plague in Madagascar during the second half of the 20th century: a survey of 20 900 notified cases. Tropical Medicine & International Health. Wiley/Blackwell (10.1111); 2006;11:1228–37.

14. Rahelinirina S, Rajerison M, Telfer S, Savin C, Carniel E, Duplantier J-M. The Asian house shrew Suncus murinus as a reservoir and source of human outbreaks of plague in Madagascar. Vinetz JM, editor. PLoS Negl Trop Dis. Public Library of Science; 2017;11:e0006072.

15. Link VB. Plague on the high seas. Public Health Rep. 1951;66:1466–72.

16. Sebbane F, Jarrett CO, Gardner D, Long D, Hinnebusch BJ. Role of the Yersinia pestis plasminogen activator in the incidence of distinct septicemic and bubonic forms of flea-borne plague. Proceedings of the National Academy of Sciences. National Acad Sciences; 2006;103:5526–30.

17. Neefjes J, Jongsma MLM, Paul P, Bakke O. Towards a systems understanding of MHC class I and MHC class II antigen presentation. Nat Rev Immunol. Nature Publishing Group; 2011;11:823–36.

18. Murphy K, Weaver C. Janeway’s Immunobiology, 9th edition. Garland Science; 2016.

19. Sironi M, Cagliani R, Forni D, Clerici M. Evolutionary insights into host-pathogen interactions from mammalian sequence data. Nature Publishing Group. Nature Publishing Group; 2015;16:224–36.

20. Brockhurst MA, Chapman T, King KC, Mank JE, Paterson S, Hurst GDD. Running with the Red Queen: the role of biotic conflicts in evolution. Proc. Biol. Sci. 2014;281:20141382–2.

21. Dyer MD, Neff C, Dufford M, Rivera CG, Shattuck D, Bassaganya-Riera J, et al. The human-bacterial pathogen protein interaction networks of Bacillus anthracis, Francisella tularensis, and Yersinia pestis. Rénia L, editor. PLoS ONE. Public Library of Science; 2010;5:e12089.

22. Chung LK, Bliska JB. Yersinia versus host immunity: how a pathogen evades or triggers a protective response. Current Opinion in Microbiology. 2016;29:56–62.

23. Shannon JG, Hasenkrug AM, Dorward DW, Nair V, Carmody AB, Hinnebusch BJ. Yersinia pestis Subverts the Dermal Neutrophil Response in a Mouse Model of Bubonic Plague. mBio. 2013;4:e00170–13–e00170–13.

24. Shannon JG, Bosio CF, Hinnebusch BJ. Dermal Neutrophil, Macrophage and Dendritic Cell Responses to Yersinia pestis Transmitted by Fleas. Monack DM, editor. PLoS Pathog. 2015;11:e1004734.

25. Gonzalez RJ, Lane MC, Wagner NJ, Weening EH, Miller VL. Dissemination of a Highly Virulent Pathogen: Tracking The Early Events That Define Infection. Valdivia RH, editor. PLoS Pathog. 2015;11:e1004587.

26. Nham T, Filali S, Danne C, Derbise A, Carniel E. Imaging of bubonic plague dynamics by in vivo tracking of bioluminescent Yersinia pestis. PLoS ONE. 2012;7:e34714.

27. Yang H, Wang T, Tian G, Zhang Q, Wu X, Xin Y, et al. Host transcriptomic responses to pneumonic plague reveal that Yersinia pestis inhibits both the initial adaptive and innate immune responses in mice. Int. J. Med. Microbiol. 2017;307:64–74.

28. Comer JE, Sturdevant DE, Carmody AB, Virtaneva K, Gardner D, Long D, et al. Transcriptomic and innate immune responses to Yersinia pestis in the lymph node during bubonic plague. Infection and Immunity. American Society for Microbiology Journals; 2010;78:5086–98.

29. Begon M, Klassovskiy N, Ageyev V, Suleimenov B, Atshabar B, Bennett M. Epizootiologic Parameters for Plague in Kazakhstan. Emerging Infect. Dis. Centers for Disease Control and Prevention; 2006;12:268–73.

30. Zhang Y, Dai X, Wang X, Maituohuti A, Cui Y, Rehemu A, et al. Dynamics of Yersinia pestis and its antibody response in great gerbils (Rhombomys opimus) by subcutaneous infection. PLoS ONE. 2012;7:e46820.

31. Casanova J-L, Abel L. The genetic theory of infectious diseases: a brief history and selected illustrations. Annu Rev Genomics Hum Genet. Annual Reviews; 2013;14:215–43.

32. Tollenaere C, Rahalison L, Ranjalahy M, Rahelinirina S, Duplantier JM, Brouat C. CCR5 polymorphism and plague resistance in natural populations of the black rat in Madagascar. Infection, Genetics and Evolution. Elsevier; 2008;8:891–7.

33. Blanchet C, Jaubert J, Carniel E, Fayolle C, Milon G, Szatanik M, et al. Mus spretus SEG& sol Pas mice resist virulent Yersinia pestis, under multigenic control. Genes and Immunity. Nature Publishing Group; 2010;12:23–30.

34. Busch JD, Van Andel R, Cordova J, Colman RE, Keim P, Rocke TE, et al. Population differences in host immune factors may influence survival of Gunnison’s prairie dogs (Cynomys gunnisoni) during plague outbreaks. Journal of Wildlife Diseases. 2011;47:968–73.

35. Demeure CE, Blanchet C, Fitting C, Fayolle C, Khun H, Szatanik M, et al. Early Systemic Bacterial Dissemination and a Rapid Innate Immune Response Characterize Genetic Resistance to Plague of SEG Mice. Journal of Infectious Diseases. 2011;205:134–43.

36. Vladimer GI, Weng D, Paquette SWM, Vanaja SK, Rathinam VAK, Aune MH, et al. The NLRP12 inflammasome recognizes Yersinia pestis. Immunity. 2012;37:96–107.

37. Tollenaere C, Jacquet S, Ivanova S, Loiseau A, Duplantier JM, Streiff R, et al. Beyond an AFL. genome scan towards the identification of immune genes involved in plague resistance in Rattus rattus from Madagascar. Mol Ecol. Wiley/Blackwell (10.1111); 2012;22:354–67.

38. Busch JD, Van Andel R, Stone NE, Cobble KR, Nottingham R, Lee J, et al. The innate immune response may be important for surviving plague in wild gunnison’s prairie dogs. Journal of Wildlife Diseases. 2013;49:920–31.

39. Tollenaere C, Ivanova S, Duplantier J-M, Loiseau A, Rahalison L, Rahelinirina S, et al. Contrasted Patterns of Selection on MHC-Linked Microsatellites in Natural Populations of the Malagasy Plague Reservoir. Salamin N, editor. PLoS ONE. Public Library of Science; 2012;7:e32814.

40. Cobble KR, Califf KJ, Stone NE, Shuey MM, Birdsell DN, Colman RE, et al. Genetic variation at the MHC DRB1 locus is similar across Gunnison’s prairie dog (Cynomys gunnisoni) colonies regardless of plague history. Ecol Evol. 2016;6:2624–51.

41. Bean AGD, Baker ML, Stewart CR, Cowled C, Deffrasnes C, Wang L-F, et al. Studying immunity to zoonotic diseases in the natural host - keeping it real. Nat Rev Immunol. Nature Publishing Group; 2013;13:851–61.

42. Monzón-Casanova E, Rudolf R, Starick L, Müller I, Söllner C, Müller N, et al. The Forgotten: Identification and Functional Characterization of MHC Class II Molecules H2-Eb2 and RT1-Db2. J. Immunol. American Association of Immunologists; 2016;196:988–99.

43. Ballingall KT, Bontrop RE, Ellis SA, Grimholt U, Hammond JA, Ho C-S, et al. Comparative MHC nomenclature: report from the ISAG/IUIS-VIC committee 2018. Immunogenetics. Springer Berlin Heidelberg; 2018;46:333–8.

44. Braunstein NS, Germain RN. The mouse E beta 2 gene: a class II MHC beta gene with limited intraspecies polymorphism and an unusual pattern of transcription. EMBO J. European Molecular Biology Organization; 1986;5:2469–76.

45. Kumar S, Stecher G, Tamura K. MEGA7: Molecular Evolutionary Genetics Analysis Version 7.0 for Bigger Datasets. Mol. Biol. Evol. 2016;33:1870–4.

46. Waterhouse AM, Procter JB, Martin DMA, Clamp M, Barton GJ. Jalview Version 2—a multiple sequence alignment editor and analysis workbench. Bioinformatics. 2009;25:1189–91.

47. Jensen KK, Andreatta M, Marcatili P, Buus S, Greenbaum JA, Yan Z, et al. Improved methods for predicting peptide binding affinity to MHC class II molecules. Immunology. Wiley/Blackwell (10.1111); 2018;54:159.

48. Hargreaves AD, Zhou L, Christensen J, Marletaz F, Liu S, Li F, et al. Genome sequence of a diabetes-prone desert rodent reveals a mutation hotspot around the ParaHox gene cluster. 2016;:1–10.

49. Kawai T, Akira S. The role of pattern-recognition receptors in innate immunity: update on Toll-like receptors. Nat. Immunol. Nature Publishing Group; 2010;11:373–84.

50. Salazar Gonzalez RM, Shehata H, O’Connell MJ, Yang Y, Moreno-Fernandez ME, Chougnet CA, et al. Toxoplasma gondii-derived profilin triggers human toll-like receptor 5-dependent cytokine production. JIN. Karger Publishers; 2014;6:685–94.

51. Roach JC, Glusman G, Rowen L, Kaur A, Purcell MK, Smith KD, et al. The evolution of vertebrate Toll-like receptors. Proceedings of the National Academy of Sciences. 2005;102:9577–82.

52. Temperley ND, Berlin S, Paton IR, Griffin DK, Burt DW. Evolution of the chicken Toll-like receptor gene family: A story of gene gain and gene loss. BMC Genomics 2015 16:1. BioMed Central; 2008;9:62.

53. Solbakken MH, Tørresen OK, Nederbragt AJ, Seppola M, Gregers TF, Jakobsen KS, et al. Evolutionary redesign of the Atlantic cod (Gadus morhua L.) Toll-like receptor repertoire by gene losses and expansions. Sci Rep. Nature Publishing Group; 2016;6:39.

54. Barreiro LB, Ben-Ali M, Quach H, Laval G, Patin E, Pickrell JK, et al. Evolutionary Dynamics of Human Toll-Like Receptors and Their Different Contributions to Host Defense. McVean G, editor. PLOS Genetics. Public Library of Science; 2009;5:e1000562.

55. Babik W, Dudek K, Fijarczyk A, Pabijan M, Stuglik M, Szkotak R, et al. Constraint and Adaptation in newt Toll-Like Receptor Genes. Genome Biology and Evolution. 2014;7:81–95.

56. Mathur R, Oh H, Zhang D, Park S-G, Seo J, Koblansky A, et al. A Mouse Model of Salmonella Typhi Infection. Cell. Cell Press; 2012;151:590–602.

57. Oldenburg M, Krüger A, Ferstl R, Kaufmann A, Nees G, Sigmund A, et al. TLR13 Recognizes Bacterial 23S rRNA Devoid of Erythromycin Resistance–Forming Modification. Science. American Association for the Advancement of Science; 2012;337:1111–5.

58. Chevret P, Dobigny G. Systematics and evolution of the subfamily Gerbillinae (Mammalia, Rodentia, Muridae). Molecular Phylogenetics and Evolution. Academic Press; 2005;35:674–88.

59. Dhariwala MO, Olson RM, Anderson DM. Induction of Type I Interferon through a Noncanonical Toll-Like Receptor 7 Pathway during Yersinia pestis Infection. Bäumler AJ, editor. Infection and Immunity. 2017;85.

60. Amemiya K, Meyers JL, Rogers TE, Fast RL, Bassett AD, Worsham PL, et al. CpG oligodeoxynucleotides augment the murine immune response to the Yersinia pestis F1-V vaccine in bubonic and pneumonic models of plague. Vaccine. 2009;27:2220–9.

61. Saikh KU, Kissner TL, Sultana A, Ruthel G, Ulrich RG. Human monocytes infected with Yersinia pestis express cell surface TLR9 and differentiate into dendritic cells. J. Immunol. 2004;173:7426–34.

62. Celhar T, Pereira-Lopes S, Thornhill SI, Lee HY, Dhillon MK, Poidinger M, et al. TLR7 and TLR9 ligands regulate antigen presentation by macrophages. Int. Immunol. 2016;28:223–32.

63. Raetz CRH, Reynolds CM, Trent MS, Bishop RE. Lipid A modification systems in gram-negative bacteria. Annu. Rev. Biochem. Annual Reviews; 2007;76:295–329.

64. Rebeil R, Ernst RK, Gowen BB, Miller SI, Hinnebusch BJ. Variation in lipid A structure in the pathogenic yersiniae. Mol. Microbiol. Wiley/Blackwell (10.1111); 2004;52:1363–73.

65. Steimle A, Autenrieth IB, Frick J-S. Structure and function: Lipid A modifications in commensals and pathogens. Int. J. Med. Microbiol. 2016;306:290–301.

66. Foster SL, Medzhitov R. Gene-specific control of the TLR-induced inflammatory response. Clin. Immunol. 2009;130:7–15.

67. Nei M, Gu X, Sitnikova T. Evolution by the birth-and-death process in multigene families of the vertebrate immune system. Proceedings of the National Academy of Sciences. National Academy of Sciences; 1997;94:7799–806.

68. Amadou C, Younger RM, Sims S, Matthews LH, Rogers J, Kumanovics A, et al. Co-duplication of olfactory receptor and MHC class I genes in the mouse major histocompatibility complex. Hum. Mol. Genet. 2003;12:3025–40.

69. Ohtsuka M, Inoko H, Kulski JK, Yoshimura S. Major histocompatibility complex (Mhc) class Ib gene duplications, organization and expression patterns in mouse strain C57BL/6. BMC Genomics 2015 16:1. BioMed Central; 2008;9:178.

70. Hughes AL, Nei M. Evolutionary relationships of class II major-histocompatibility-complex genes in mammals. Mol. Biol. Evol. 1990;7:491–514.

71. Viluma A, Mikko S, Hahn D, Skow L, Andersson G, Bergström TF. Genomic structure of the horse major histocompatibility complex class II region resolved using PacBio long-read sequencing technology. Sci Rep. Nature Publishing Group; 2017;7:45518.

72. Katju V, Lynch M. The structure and early evolution of recently arisen gene duplicates in the Caenorhabditis elegans genome. Genetics. Genetics Society of America; 2003;165:1793–803.

73. Gregers TF, Fleckenstein B, Vartdal F, Roepstorff P, Bakke O, Sandlie I. MHC class II loading of high or low affinity peptides directed by Ii/peptide fusion constructs: implications for T cell activation. Int. Immunol. 2003;15:1291–9.

74. Liu X, Zhan Z, Li D, Xu L, Ma F, Zhang P, et al. Intracellular MHC class II molecules promote TLR-triggered innate immune responses by maintaining activation of the kinase Btk. Nat. Immunol. Nature Publishing Group; 2011;12:416–24.

75. Matzaraki V, Kumar V, Wijmenga C, Zhernakova A. The MHC locus and genetic susceptibility to autoimmune and infectious diseases. Genome Biology. BioMed Central; 2017;18:76.

76. Zhang Y, Dai X, Wang Q, Chen H, Meng W, Wu K, et al. Transmission efficiency of the plague pathogen (Y. pestis) by the flea, Xenopsylla skrjabini, to mice and great gerbils. Parasit Vectors. BioMed Central; 2015;8:256.

77. Martin M. Cutadapt removes adapter sequences from high-throughput sequencing reads. EMBnet.journal. 2011;17:pp.10–2.

78. Gnerre S, Maccallum I, Przybylski D, Ribeiro FJ, Burton JN, Walker BJ, et al. High-quality draft assemblies of mammalian genomes from massively parallel sequence data. Proc. Natl. Acad. Sci. U.S.A. National Acad Sciences; 2011;108:1513–8.

79. Elgvin TO, Trier CN, Tørresen OK, Hagen IJ, Lien S, Nederbragt AJ, et al. The genomic mosaicism of hybrid speciation. Sci Adv. 2017;3:e1602996.

80. Pujolar JM, Dalén L, Olsen RA, Hansen MM, Madsen J. First de novo whole genome sequencing and assembly of the pink-footed goose. Genomics. 2018;110:75–9.

81. Parra G, Bradnam K, Korf I. CEGMA: a pipeline to accurately annotate core genes in eukaryotic genomes. Bioinformatics. 2007;23:1061–7.

82. Parra G, Bradnam K, Ning Z, Keane T, Korf I. Assessing the gene space in draft genomes. Nucleic Acids Res. 2009;37:289–97.

83. Simão FA, Waterhouse RM, Ioannidis P, Kriventseva EV, Zdobnov EM. BUSCO: assessing genome assembly and annotation completeness with single-copy orthologs. Bioinformatics. Oxford University Press; 2015;31:3210–2.

84. Kumar S, Jones M, Koutsovoulos G, Clarke M, Blaxter M. Blobology: exploring raw genome data for contaminants, symbionts and parasites using taxon-annotated GC-coverage plots. Front Genet. Frontiers; 2013;4:237.

85. Holt C, Yandell M. MAKER2: an annotation pipeline and genome-database management tool for second-generation genome projects. BMC Bioinformatics. 2011.

86. Varadharajan S, Sandve SR, Gillard GB, Tørresen OK, Mulugeta TD, Hvidsten TR, et al. The grayling genome reveals selection on gene expression regulation after whole genome duplication. Genome Biology and Evolution. 2018.

87. Lomsadze A, Ter-Hovhannisyan V, Chernoff YO, Borodovsky M. Gene identification in novel eukaryotic genomes by self-training algorithm. Nucleic Acids Res. 2005;33:6494–506.

88. Tørresen OK, Brieuc MSO, Solbakken MH, Sørhus E, Nederbragt AJ, Jakobsen KS, et al. Genomic architecture of haddock (Melanogrammus aeglefinus) shows expansions of innate immune genes and short tandem repeats. BMC Genomics 2015 16:1. BioMed Central; 2018;19:240.

89. Hurt P, Walter L, Sudbrak R, Klages S, Müller I, Shiina T, et al. The genomic sequence and comparative analysis of the rat major histocompatibility complex. Genome Res. Cold Spring Harbor Lab; 2004;14:631–9.

90. Katoh K, Standley DM. MAFFT multiple sequence alignment software version 7: improvements in performance and usability. Mol. Biol. Evol. 2013;30:772–80.

91. Maddison WP, Maddison DR. Mesquite: a modular system for evolutionary analysis. Version 3.4.

92. Talavera G, Castresana J, Kjer K, Page R, Sullivan J. Improvement of Phylogenies after Removing Divergent and Ambiguously Aligned Blocks from Protein Sequence Alignments. Kjer K, Page R, Sullivan J, editors. Systematic Biology. Oxford University Press; 2007;56:564–77.

93. Stamatakis A. RAxML-VI-HPC: maximum likelihood-based phylogenetic analyses with thousands of taxa and mixed models. Bioinformatics. Oxford University Press; 2006;22:2688–90.

94. Aberer AJ, Krompass D, Stamatakis A. Pruning rogue taxa improves phylogenetic accuracy: an efficient algorithm and webservice. Systematic Biology. 2013;62:162–6.

95. Huelsenbeck JP, Ronquist F. MRBAYES: Bayesian inference of phylogenetic trees. Bioinformatics. 2001;17:754–5.

96. Delport W, Poon AFY, Frost SDW, Kosakovsky Pond SL. Datamonkey 2010: a suite of phylogenetic analysis tools for evolutionary biology. Bioinformatics. Oxford University Press; 2010;26:2455–7.

97. Weaver S, Shank SD, Spielman SJ, Li M, Muse SV, Kosakovsky Pond SL. Datamonkey 2.0: a modern web application for characterizing selective and other evolutionary processes. Mol. Biol. Evol. 2018;83:8916.

98. Kosakovsky Pond SL, Murrell B, Fourment M, Frost SDW, Delport W, Scheffler K. A Random Effects Branch-Site Model for Detecting Episodic Diversifying Selection. Mol. Biol. Evol. 2011;28:3033–43.

99. Murrell B, Wertheim JO, Moola S, Weighill T, Scheffler K, Kosakovsky Pond SL. Detecting individual sites subject to episodic diversifying selection. Malik HS, editor. PLOS Genetics. 2012;8:e1002764.

100. Smith MD, Wertheim JO, Weaver S, Murrell B, Scheffler K, Kosakovsky Pond SL. Less is more: an adaptive branch-site random effects model for efficient detection of episodic diversifying selection. Mol. Biol. Evol. 2015;32:1342–53.

101. Kelley LA, Mezulis S, Yates CM, Wass MN, Sternberg MJE. The Phyre2 web portal for protein modeling, prediction and analysis. Nat Protoc. Nature Publishing Group; 2015;10:845–58.

102. Péléraux A, Karlsson L, Chambers J, Peterson PA. Genomic organization of a mouse MHC class II region including the H2-M and Lmp2 loci. Immunogenetics. 1996;43:204–14.

103. Chomczynski P, Sacchi N. The single-step method of RNA isolation by acid guanidinium thiocyanate-phenol-chloroform extraction: twenty-something years on. Nat Protoc. Nature Publishing Group; 2006;1:581–5.

